# RetroMol: Parsing a shared encoding from natural products and their biosynthetic gene clusters

**DOI:** 10.64898/2026.06.12.731935

**Authors:** David Meijer, Sam E. Williams, Barbara Terlouw, Pep Charusanti, Leron Kok, Michael A. Skinnider, Tilmann Weber, Justin J. J. van der Hooft, Alan R. Healy, Marnix H. Medema

**Affiliations:** Bioinformatics Group, Wageningen University & Research, Wageningen, the Netherlands; Department of Biotechnology and Biomedicine, Technical University of Denmark, Lyngby, Denmark; Princess Máxima Center for Pediatric Oncology, Utrecht, the Netherlands; Oncode Institute, Utrecht, the Netherlands; Lewis-Sigler Institute for Integrative Genomics, Princeton University, Princeton, the United States of America; Ludwig Institute for Cancer Research, Princeton Branch, Princeton, NJ, USA; Department of Biochemistry, University of Johannesburg, Johannesburg, South Africa; Chemistry Program, New York University Abu Dhabi, Saadiyat Island, Abu Dhabi, United Arab Emirates

## Abstract

Natural products such as polyketides and nonribosomal peptides (NRPs) are important sources of bioactive compounds, including many antibiotics. Many of them are assembled by modular enzyme complexes and further modified and diversified by tailoring reactions encoded by biosynthetic gene clusters (BGCs). Although natural products and their coding BGCs describe different data modalities of the same biochemical process, a unified language to jointly describe their biochemistry is lacking. Here we introduce a sequence-based representation of the core biosynthesis of modular natural products, which we call primary sequences, that bridges chemical structures and BGCs. We also present RetroMol, an algorithm that parses either natural product structures or their encoding BGCs into their primary sequences of natural product building blocks. RetroMol allows for similarity scoring between natural products and BGCs, enabling the retrieval of compounds, BGCs, and a combination of the two, based on their biosynthetic similarity. This can, for instance, be used to retrieve biosynthetically similar but structurally dissimilar compounds, or link natural products to candidate coding BGCs in large experimental datasets. We demonstrate the latter by rediscovering the nocardichelin B BGC as a proof of principle. We also exemplify the utility of biosynthetic similarity by showing various pairs of biosynthetically similar compounds with low structural similarity. Together, these results establish primary sequences as a shared biosynthetic encoding for natural product comparison and BGC prioritization.

## Introduction

Bacteria and fungi make specialized metabolites, or natural products, that help them survive and compete in their environments. These compounds are usually produced by biosynthetic gene clusters (BGCs) and are made only when beneficial, since their synthesis can be metabolically expensive^1^. Important natural products classes include polyketides, nonribosomal peptides, and hybrid molecules, which are built iteratively from small biochemical units by modular enzymatic assembly lines^2–4^. Additional cyclization events and post-assembly modifications then expand their structural and functional diversity^5^. Given that this process has generated numerous bioactive compounds, including clinically important antibiotics, microbial natural products represent a valuable source for drug discovery and biotechnology^6^.

In parallel with advances in genome mining and structure curation, the amount of data available for natural product discovery has grown rapidly. antiSMASH^7,8^ now enables large-scale detection and annotation of BGCs, and public resources contain hundreds of thousands of mined microbial gene clusters. Moreover, NPAtlas^9^ and other databases catalog tens of thousands of curated microbial natural product structures.

Several specialized frameworks have been developed in recent years that link modular BGCs to their corresponding natural products. Nerpa 2^10^ focuses exclusively on NRPs, the BioPKS pipeline^11^ is designed for polyketides, and Seq2Saccharide^12^ was specifically designed for saccharide natural products. The PRISM suite^13^, including GRAPE^14^, also maps BGCs to modular natural products. CHAMOIS^15^ employs multiple logistic regression classifiers to associate structural features with Pfam domains. An early proof-of-concept for a metabolite-to-BGC strategy was provided by Khater et al.^16^, describing a retrobiosynthetic approach that links metabolite functional groups to biochemical transformations. Methods for comparing BGCs, like BiG-SCAPE 2^17^, also exist. Comparing compounds is mainly done through comparing calculated or learned structural fingerprints^18^.

Although many methods exist for comparing compounds with BGCs, BGCs with BGCs, and compounds with compounds, a shared encoding for describing their biochemistry jointly is still missing. Here we introduce a representation that jointly describes the core biosynthesis of modular natural products and their coding BGCs as sequences of biosynthetic building blocks (Figure 1), which we call primary sequences. We have also developed an algorithm called RetroMol that efficiently decomposes natural products structures and antiSMASH-annotated BGCs into primary sequences. We further show that the same representations provide a more biosynthetically meaningful measure of compound relatedness than conventional chemical similarity metrics. Finally, we demonstrate that RetroMol-mined primary sequences can link natural products to their coding BGCs in large experimental datasets, rediscovering the nocardichelin B BGC as a proof of principle.

**Figure 1:**
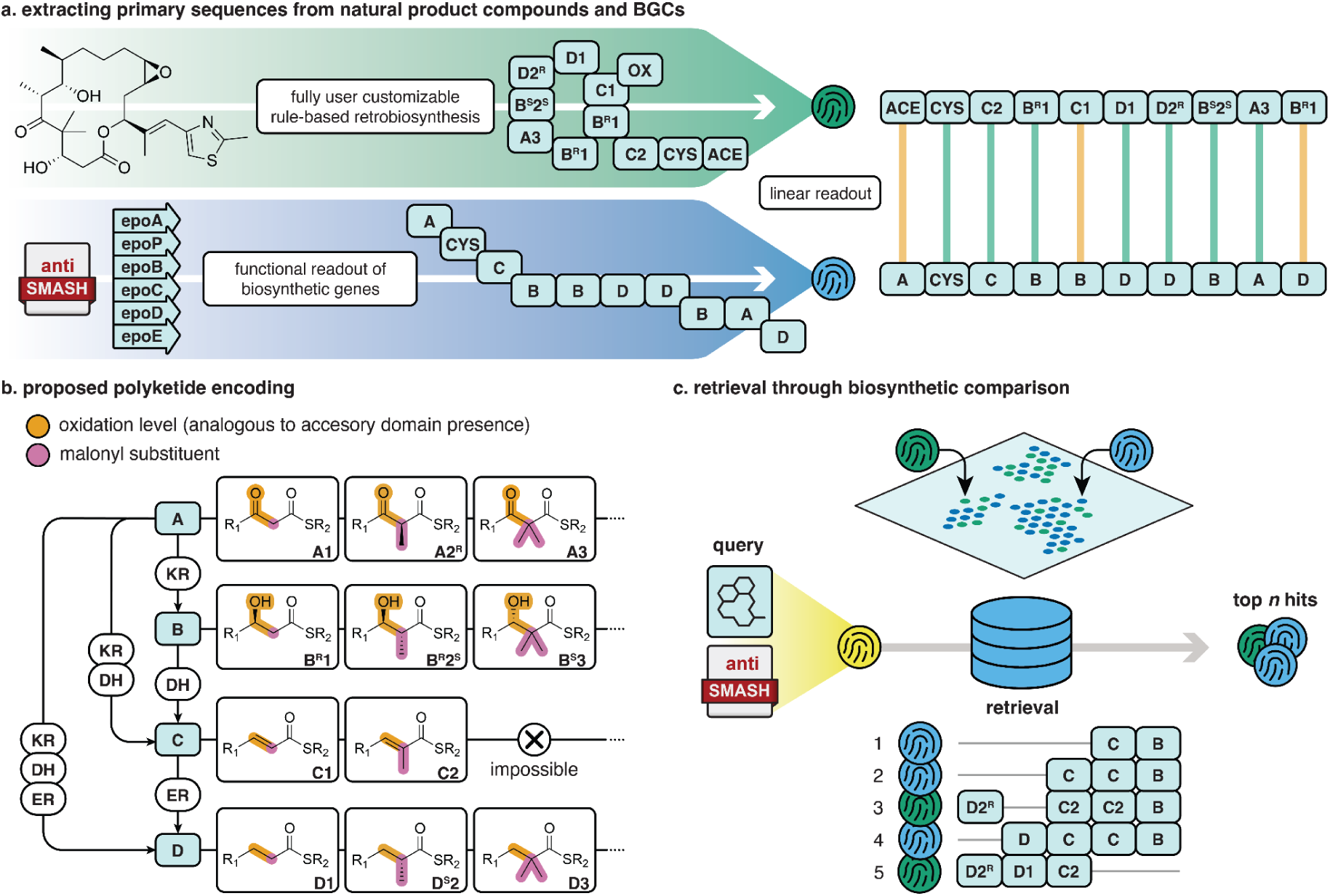
Overview of the RetroMol algorithm. (a) Modular natural products (top) and BGCs (bottom) are featurized into primary sequences. (b) Our proposed polyketide encoding to featurize polyketide structures is shown, analogous to polyketide biosynthesis. Other modular building blocks like amino acids are represented by the first unique set of three alphanumeric characters from their name. Proteinogenic amino acids are assigned their common three-letter abbreviation. (c) Primary sequence space can be queried through alignment of raw sequences or by comparing primary sequence embeddings to retrieve biosynthetically similar compounds and BGCs.

## Results

The RetroMol algorithm encodes natural product compounds and BGCs into a shared primary sequence space. The sequence elements are cross-modal, meaning that they describe structural features that can be inferred either from chemical structures or from BGCs annotated by antiSMASH. Primary sequences not only enable the retrieval of similar compounds with a compound, or similar BGCs with a BGC, i.e., uni-modal, but also of compounds with a BGC and BGCs with a compound, i.e., cross-modal. The cross-modal functionality of RetroMol also enables annotation transfer across modalities: for instance, one can infer BGC-associated information such as the taxonomy of a producing organism directly from structure. The reverse is likewise true: one can use a BGC to query compound space and retrieve typical compound-associated information like experimentally validated biological activities.

### A common biosynthetic language encoding for modular natural products and their coding BGCs

Modular natural products such as type I polyketides, nonribosomal peptides (NRPs), and polyketide-NRP hybrids can be interpreted through a retrosynthetic, biosynthesis-aware lens. In their biosynthetic logic, these molecules originate as linear sequences of biochemical building blocks (polyketide extender units and/or amino-acid-derived monomers) that may cyclize and acquire tailoring modifications, producing the final structure. RetroMol leverages this logic by deconstructing a compound structure into an ordered primary sequence of building blocks using custom retrobiosynthetic rules. This linear representation of building blocks can then readily be compared across related structures, while retaining a high degree of biosynthetic interpretability.

Representing natural product structures as primary sequences requires a standardized vocabulary for biosynthetic building blocks. For NRPs, monomers can be expressed directly using conventional names (e.g., proteinogenic amino acids and common nonproteinogenic variants). For polyketides, in contrast, extender-derived units lack a consistent naming convention. We therefore introduce a structured polyketide encoding that standardizes extender-derived units along two biosynthetically meaningful axes: the β-carbon oxidation state and the ɑ-carbon substituent identity. The β-carbon oxidation state reflects the ketoreductase (KR), dehydratase (DH), and enoylreductase (ER) reduction series, while the digit code reflects extender choice (e.g., malonyl-vs methylmalonyl-CoA-derived monomers). This encoding yields tokens that mirror how PKS modules specify chemistry and support controlled coarsening when only partial information is available. The logic is shown in Figure 1b, and the full scheme is provided in Figure S1. Additionally, the chirality of either carbon atom can be annotated with either *R*/*S*.

Having defined an “alphabet” for the representation of natural product structures, we populate the same token space from BGCs to enable direct comparison between modalities. Although RetroMol is extensible to other natural product classes, we focus here on type I cis-and trans-AT PKSs and NRPSs. In NRPS BGCs, each module’s adenylation (A) domain activates a monomer for incorporation. We predict the substrate of these A domains using PARAS^19^, a random forest model trained experimentally validated substrates spanning proteinogenic, nonproteinogenic, and specialized building blocks. For polyketide BGCs, we use antiSMASH^7,8^ domain calls to extract ketosynthase (KS) domains and use them to anchor a polyketide module. Then, we match these KS anchors to their nearest downstream acyltransferase (cis-AT) domain or an upstream AT domain (trans-AT). Presence of ketoreductase (KR), dehydratase (DH), and enoylreductase (ER) accessory domains are then used to assign the β-carbon letter in our encoding. The order of modules within an antiSMASH identified candidate cluster defines the primary sequence. As exemplified by the primary sequences extracted from the epothilone B structure and its coding BGCs, primary sequences don’t always exactly line up (Figure 1a). Late stage modifications of the structure after biosynthesis of the core structure are reflected in the structure, but cannot be reliably predicted from the BGC alone. Nevertheless there is enough overlap to facilitate a reasonable match.

### A customizable compound parser with high structural coverage for polyketides and peptides

To illustrate RetroMol’s retrobiosynthetic parsing performance, we applied it to 36,454 microbial natural products from NPAtlas version 3.0 (Figure 2a). RetroMol parsing quality is assessed using coverage, defined as the fraction of heavy atoms in a compound that are mapped to a monomer. A coverage of ≥0.9 indicates that the default ruleset (S4-5) successfully parsed the compound to a high degree and that nearly all monomers were identified. A coverage of ≥0.5 suggests that at least half of the compound was parsed. 12,788 compounds (35%) have a coverage of at least 0.5, while 8,001 compounds (22%) have a coverage of at least 0.9. Top parseable compounds are generally classified as peptides and polyketides by NPClassifier^2^.

**Figure 2:**
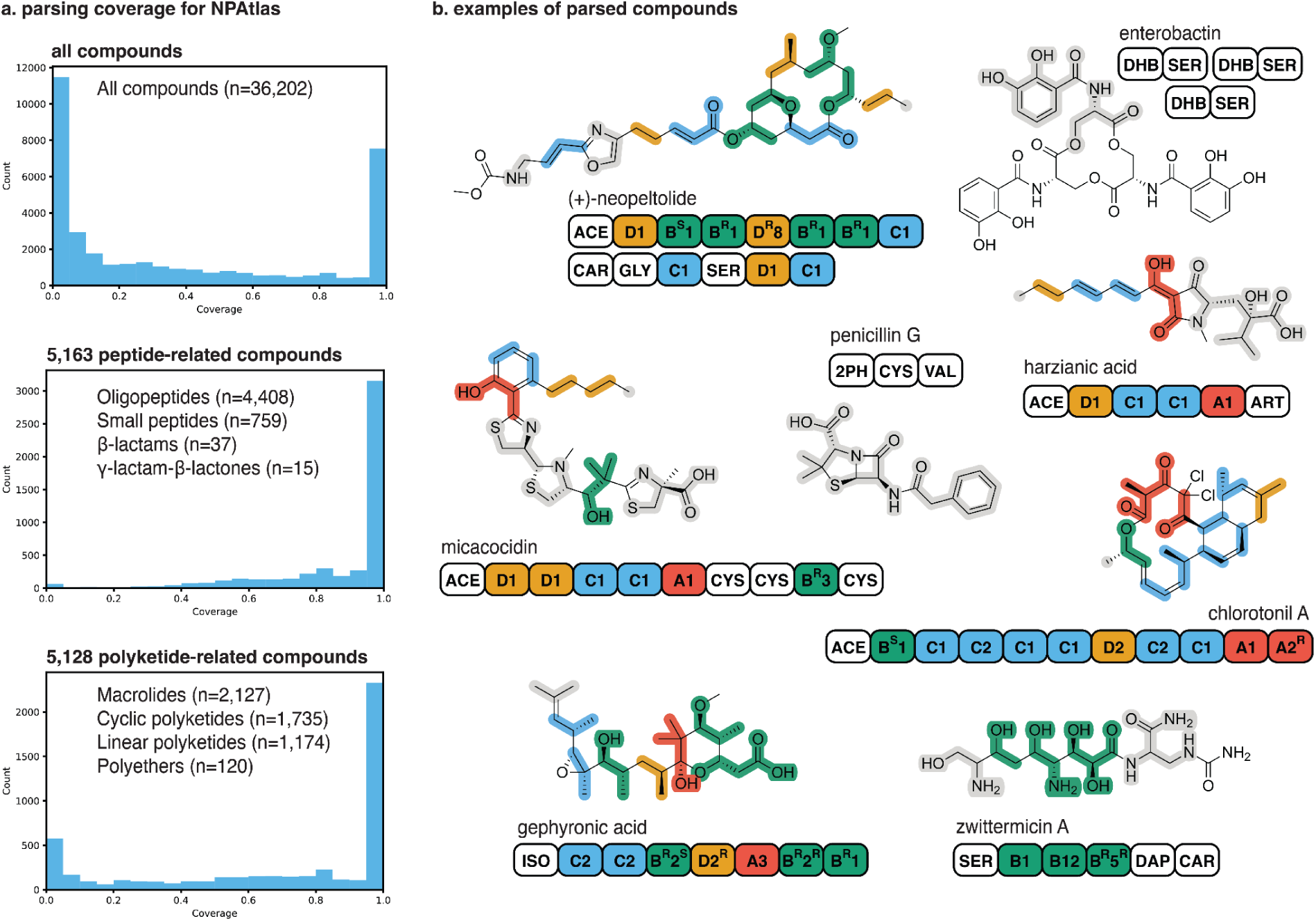
Parsing capabilities of the RetroMol parsing algorithm. (a) Parsing coverage for all NPAtlas compounds, for polyketide-related classes, and for peptide-related classes. (b) Examples of different compounds parsed by RetroMol.

Coverage of specific subclasses of natural products is high as primary sequences are designed to describe modular natural products. For example, of 5,163 peptide-related compounds, 3,428 had a ≥0.9 coverage (66%). Of 5,128 compounds classified to be a polyketide, 2,436 had a ≥ 0.9 coverage (48%). Furthermore, 14,555 of all compounds (40%) have a coverage of at most 0.1, whilst for peptides this was 75 compounds (0.01%) and for polyketides this was 759 (15%). This indicates that most poorly parsable compounds with the default rule set are unlikely to be type I polyketides or NRPs. The default ruleset can be supplemented with reaction and monomer rules to parse other natural product classes that currently cannot be parsed.

Example retrobiosyntheses are shown in Figure 2b. RetroMol can separate tailoring moieties such as acetylations (e.g., (+)-neopeltolide), oxidations (e.g., gephyronic acid), halogenations (e.g., chlorotonil A), and methylations (e.g., gephyronic acid and harzianic acid), as well as aminations and glycosylations from core structures. Furthermore, because the algorithm only considers C-C and C-N bonds as backbone bonds, compound backbones are split up at C-O bonds (e.g., neopeltolide and enterobactin). RetroMol’s compound parsing algorithm was set up to be customizable. In the case of harzianic acid, this customizability is particularly apparent: by adding a dedicated monomer definition for its characteristic 4,4-disubstituted glutamic acid^20^, we ensure it is treated as a single protected building block and is not decomposed further by the reaction rules.

RetroMol differs fundamentally from state-of-the-art parsers used by GRAPE^14^ and Nerpa 2^10^. GRAPE outputs text-based monomer sequences but does not provide atom mappings of monomers back to the input compound, and its polyketide annotation is limited to broad categories such as malonyl or methylmalonyl with simple reduction-state tags. RetroMol instead returns a retraceable assembly graph (figure 1a top workflow), enabling explicit atom mapping and richer polyketide encodings. Nerpa 2 also uses graph-based outputs but lacks support for polyketides. Neither GRAPE nor Nerpa implement stereochemistry-aware matching, which is implemented in RetroMol and can be turned on if needed with available corresponding matching rules. Finally, CHAMOIS^15^ does not extract monomers at all; instead, it predicts binary fingerprints from BGC domains.

### Conserved bioactivity reflects shared biosynthetic patterns

Retrieval between compounds using biosynthetic fingerprints and fine-tuning through alignment prioritizes similarity in monomer composition. As a result, compounds with divergent final structures but shared biosynthetic patterning can be linked, even when structure-based fingerprints fail to do so^21^. This is illustrated by the polyketides dictyostatin and discodermolide (Figure 3a top alignment). Biosynthetically, these compounds are highly similar due to their overlap and order in biosynthetic building blocks (cosine similarity of 0.95, normalized alignment score of 0.71), whereas their Morgan structural fingerprints are less similar because of differences in cyclization (Tanimoto coefficient of 0.20). Despite these chemical differences, both compounds are potent tubulin inhibitors as they both adopt nearly identical bioactive conformations at the microtubule binding site^22,23^. This demonstrates that conserved biosynthetic architecture can sometimes preserve shared bioactivity, even when compound structures appear diverse at first glance. Nevertheless, this relationship is not universal and small structural changes can also give rise to activity cliffs.

**Figure 3:**
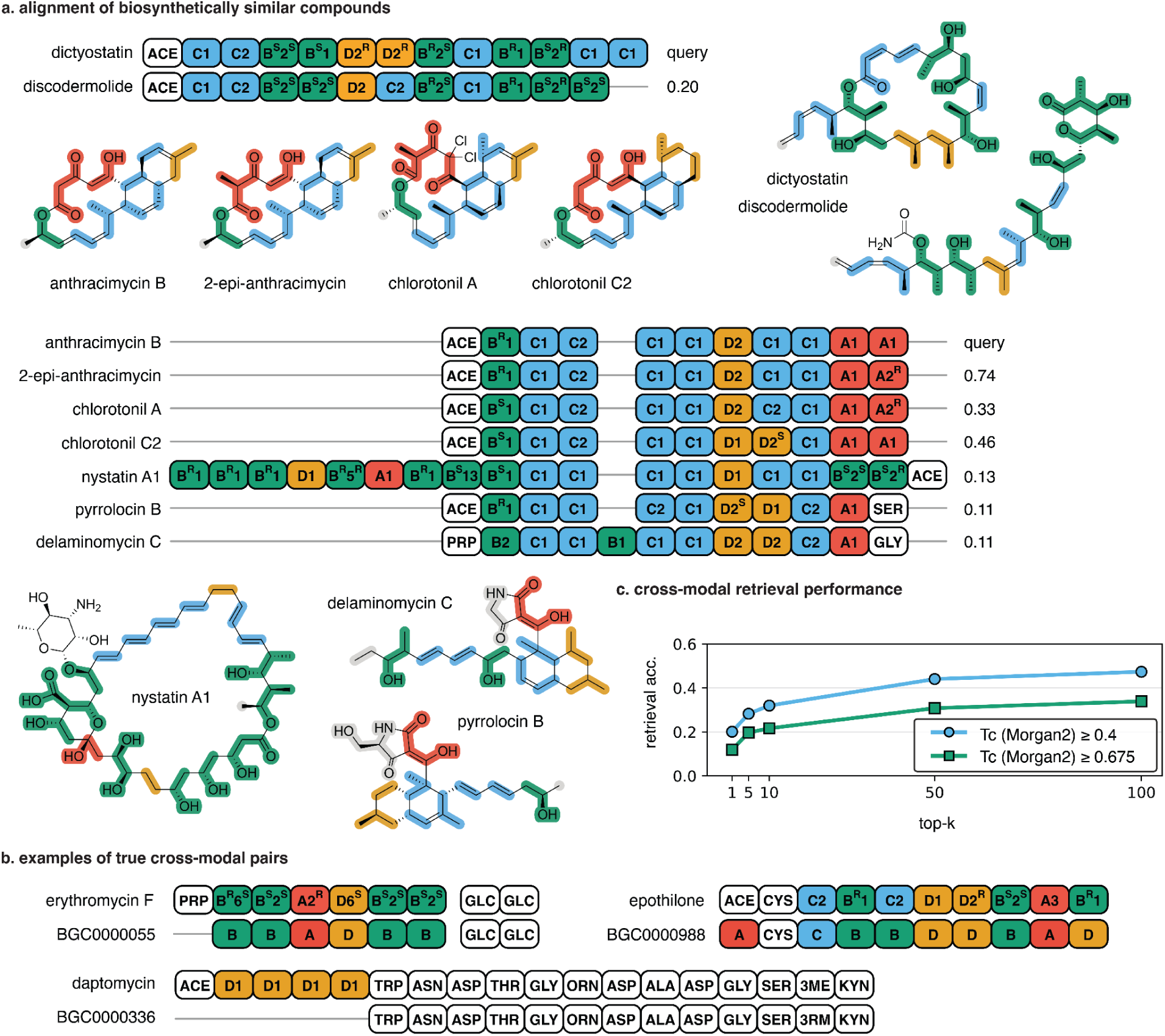
Cross-modal and unimodal performance and examples. (a) Aligned structural readouts of dictyostatin and discodermolide, and anthracimycin B and some of its nearest neighbors in modular biosynthetic space. Listed scores are pairwise Tanimoto similarities to the query, based on Morgan (radius 2, 2048 bits, considering stereochemistry) fingerprints. (b) Cross-modal alignment of erythromycin C, epothilone, and daptomycin and their corresponding BGCs. (c) Cross-modal retrieval scores for 513 MIBiG BGC-compound pairs.

Anthracimycin B provides a second example of how biosynthetic fingerprints group biosynthetically similar but structurally divergent structures (Figure 3a, bottom alignment). Anthracimycin B, 2-epi-anthracimycin, and chlorotonil structures share highly similar polyketide-derived primary sequences, consistent with a conserved assembly-line logic^24^. In contrast, pyrrolocin B and delaminomycin C retain a closely related polyketide backbone but terminate with an amino-acid-derived unit (e.g., Ser/Gly), which changes the identity and position of nucleophiles available for ring closure and thus implies a different cyclization regime than the purely polyketide structures. This example highlights how RetroMol provides insights into small edits at the level of primary sequences, such as the addition of a terminal amino-acid-like building block, which drives substantial downstream structural diversity by altering macrocyclization. Additionally, the CCD(C|D)C motif pattern seems related to the presence of a Diels-Alder-derived carbocycle, which only nystatin A1 lacks. This allows us to describe structural features in a descriptive, biosynthesis-oriented way.

Together, these examples illustrate how biosynthetic fingerprints and primary sequences provide an insightful way to identify and investigate relationships among biosynthetically similar yet structurally divergent metabolites and to describe them succinctly.

### Fingerprint-based retrieval facilitates fast and accurate mapping of metabolite-BGC similarities

Since primary sequences are a shared biosynthetic encoding between natural products and BGCs, they can be used to retrieve one with the other. Primary sequences parsed from compounds describe more detailed information than what we can currently reliably predict from BGCs, such as a polyketide motif’s exact malonyl substituent and its stereochemistry (Figure 3b). Nevertheless, there are plenty of biosynthetic signals to filter out eligible compound matches for a BGC or vice versa. This is demonstrated using 513 MIBiG-sourced NRP and type I polyketide coding BGCs to retrieve an associated compound from NPAtlas (Figure 3c)^9,25^. For 20% of BGCs, the top-1 retrieved compound has at least a 0.4 Tanimoto coefficient (Morgan 2, 2048) to the BGC’s true compound. This number rises to 47% for the first 100 compounds. A Tanimoto coefficient of 0.4 has been defined by the authors of MS2Mol, a tool that predicts compound structures from mass spectra *de novo*, as “meaningfully similar”^26^. 12% of the top-1 retrieved compounds had a Tanimoto coefficient (Morgan 2, 2048 bits) of at least 0.675, which is described by the same authors as “close match”. This shows that primary sequences contain useful biosynthetic signals to retrieve correct matches. Nevertheless, the retrieval is not perfect which can be attributed to a “noisy” biosynthetic signal or imperfect parsing. For example, the BGC for the NRP thiocoraline (BGC0000445) contains many more NRP modules than found monomers in the structure. Also, the compound might not always be (fully) parseable with the current RuleSet. Only five polyketide motifs can be parsed from the abyssomicin C structure, while its corresponding BGC0000001 reports seven biosynthetic modules.

Although there is expected overlap between primary sequences from compounds and BGCs, compound primary sequences rarely exactly match the BGC primary sequences. BGCs may incorporate modules iteratively or non-linearly, modules can have inactive domains, substrate specificity for modules might be ambiguous or inaccurately predicted, and compounds encoded by BGCs often undergo post-assembly modifications. An example is the C2 methylation in chlorotonil A (Figure 2a). Retrobiosynthesis of the chlorotonil A structure yields an A2 module, whereas the polyketide assembly line was experimentally shown to incorporate an A1 extender unit that is subsequently methylated^27^.

As BGC inference tools improve, predictions of incorporated substructures and enzymatic modifications will become more accurate. RetroMol is designed to support these advances by enabling the addition of more predictors. For example, the BGC parser uses a Pfam model to find glycosyltransferases, and emits a glycosylation family token upon identification. This enables the matching of tailoring enzymes and moieties, extending beyond primary sequences.

At present, usable BGC information is limited primarily to predicted polyketide extender units from type I PKS cis and trans AT domains using antiSMASH, and substrate specificity predictions from NRPS A domains using PARAS^19^. Because these features represent only a subset of BGC-encoded chemistry, the maximum achievable similarity between the primary sequence of a BGC and that of a highly modified compound is inherently low, even with perfect structure parsing. This is exemplified in Figure 3a. For erythromycin, RetroMol’s compound parser parses a six-extender unit backbone with a propionate starter from the structure, but the RetroMol BGC parser reliably predicts only the extender unit oxidation states. Although RetroMol parses an acetate starter unit out of the epothilone structure, the EpoA domain architecture in its coding BGCs indicates that this starter is generated from a malonyl-loaded ACP via a decarboxylative loading mechanism rather than by direct acetyl loading^28^. For daptomycin, the decanoic acid unit cannot be inferred directly from the BGC’s Condensation starter domain, while at the same time our default ruleset digests the found decanoic acid as a polyketide structure. Despite these gaps, the aligned gene-structure primary linear sequence readouts shown in Figure 3b, inspired by Pep2Path^29^, are clearly comparable. Using only the polyketide primary sequence parsed from BGC0000055, we retrieve erythromycin F as the top-1 hit from the parsed NPAtlas database, followed by other erythromycins and megalomicins (cosine similarity of 0.80, normalized alignment score of 0.97). We can potentially make the filter step with the fingerprint even more discriminating by including a Pfam model for detecting glycosyltransferases when parsing the BGC, which RetroMol allows for. Querying the parsed NPAtlas database with BGC0000988 retrieved a top-1 hit with epothilone N (cosine similarity of 0.85, normalized alignment score of 0.82), and querying the parsed NPAtlas database with BGC00000336 retrieved a top-1 hit with taromycin B (cosine similarity of 0.93, normalized alignment score of 0.68), followed by 21978C1(D-Asn11), 21978C2(D-Asn11), and 21978C3(D-Asn11). All of these compounds are close analogs of daptomycin, which is a remarkable find as daptomycin is not present in NPAtlas.

Overall, biosynthetic fingerprints and primary sequences enable useful metabolite-BGC matching, even when current genome-mining and prediction tools capture only part of the encoded biosynthetic logic.

### RetroMol biosynthetic fingerprints enable deorphanization and functional annotation of BGCs

We showcase RetroMol’s deorphanization capability by linking the known siderophore nocardichelin B^30^ to its hybrid NRPS and NRPS-independent siderophore (NIS) BGC, present in an environmental *Nocardia* strain from a large collection of sequenced actinomycetes^31^. Four environmental *Nocardia* strains were cultivated under different conditions, and the cultures analysed. Untargeted liquid chromatography coupled to tandem mass spectrometry (LC-MS/MS) data revealed a feature (716.4593 m/z) that was only observed in *Nocardia sp.* NBC_01329, with a reasonably close match to the nocardichelin B GNPS reference spectra (Modified cosine score: 0.76937, S2).

Nocardichelin B contains a tetradec-2-enoic acid (TET), two N-(5-aminopentyl)hydroxylamines (N5A) linked by a butanedioic acid (BUT), and also includes serine and salicylic acid (Figure 4a). The N5A-BUT-N5A motif is a typical pattern seen in other siderophores, such as tenacibactin and desferrioxamine, and these biosynthetic similar compounds were identified using the biosynthetic fingerprint of nocardichelin B after querying the database with only the chelating repeats and the terminal appendage (Figure 4b).

**Figure 4:**
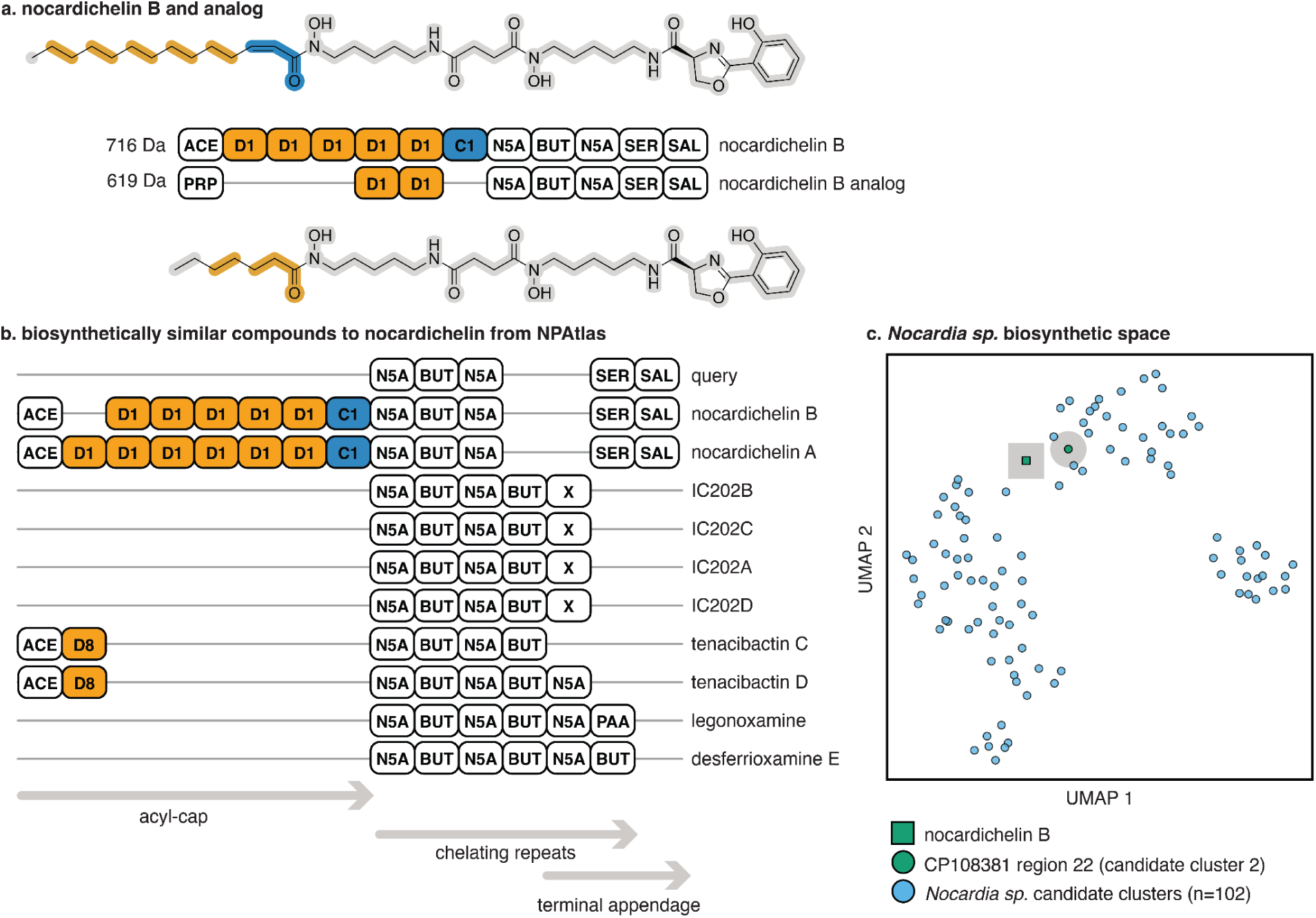
Deorphanization results. (a) Structures of nocardichelin B and its analog with highlighted monomers and aligned primary sequences. (b) Biosynthetically similar compounds to nocardichelin B. (c) Nocardichelin B structural readout co-embedded with 102 *Nocardia sp.* candidate gene cluster readouts in a UMAP of biosynthetic fingerprint space.

Nocardichelin B is a relatively challenging case for deorphanization compared to larger multimodular systems, as it only has two NRPS modules. Therefore, we manually added a *siderophore* pseudonym label to N5A and BUT monomers in RetroMol’s default dataset to help match compounds with similar chelating repeats to BGC clusters classified by antiSMASH as siderophore-producing. Although antiSMASH does not currently predict chelating repeats directly, adding the siderophore label enables RetroMol to match these compounds to BGCs more effectively. This example illustrates how RetroMol’s design, although focused on fully modular NRPs and type I polyketides, can be readily extended to other classes of natural products.

To identify the candidate BGC for nocardichelin B, we first mined the four *Nocardia* genomes with antiSMASH 7.0 and subsequently co-embedded the 136 found BGC regions (containing 102 candidate clusters with at least two modules in total), with the nocardichelin B structure using the RetroMol fingerprinter. Additionally, we added a *siderophore* label to the BGC-derived fingerprint whenever the product was predicted by antiSMASH to be a siderophore. The UMAP of this embedding space provides an intuitive and interpretable overview of similarity relationships (Figure 4c). Among all candidate clusters, BGC region 22, candidate cluster 2 from *Nocardia sp*. NBC_01329 ranked second closest in fingerprint space to the nocardichelin B fingerprint using cosine similarity alone (cosine similarity of 0.40). This association is driven by shared cross-modal features, inferred from both BGC and compound modalities, correctly identifying the BGC in the sole producing strain. The first hit (BGC region 12, candidate cluster 1 from *Nocardia sp.* NBC_01730) had a slightly higher cosine similarity of 0.45, but did not contain a single predicted amino acid substrate and was therefore discarded. RetroMol is effective at BGC prioritization, but manual inspection remains key.

Matching to the previously mentioned candidate cluster was driven primarily by the A domain substrate predictions. Notably, the match is strengthened not only by the presence of the expected substrates, but also by the absence of additional, unrelated A domain predictions: clusters encoding serine alongside many extra monomers yield substantially poorer agreement because their primary sequences introduce mismatching features. PARAS^19^ predicted the second A domain as N6-hydroxylysine rather than the structurally similar N5A-like substrate. However, the prediction confidence was low (0.436).

During the preparation of this manuscript, the BGC of nocardichelin B was identified by Fisk *et al.* in the strain *Nocardia carnea* NBRC 14403^32^. Interestingly, Fisk et al. note promiscuity in the acyl-CoA substrate-loading unit across the NcdE-ATx domain, with tolerance for shorter but not longer fatty acyl chains. In our metabolomics data, we identify a related feature (620.293 m/z) that we propose to represent a shorter-chain nocardichelin analogue (Figure 4a & S2). In support of this, the spectral comparison tool ModiFinder^33^ indicated a high likelihood that the modification was located on the acyl chain, further supporting this promiscuity, which is consistent with observations in other siderophore systems^34^. Closer examination of the reported cluster and our prioritized region showed that the clusters were highly similar, confirming we prioritized the correct cluster for nocardichelin B.

However, we observed a few differences: our cluster encodes a putative siderophore-interacting protein, homologous to ferric vibriobactin reductase ViuB, which mediates intracellular iron release from the siderophore complex^35^. Our cluster lacked homologues for *ncdA* (aryl adenylating enzyme) and *ncdB* (salicylate synthase) within the region boundaries. However, a BLAST search identified adjacent homologues of *ncdA* and *ncdB* located approximately 170 kb upstream in the genome, showing these functions can be encoded outside the biosynthetic locus. We also identified three additional ABC transporter genes (*ncdQRS*), present in both clusters but not discussed in Fisk *et al.,* which may be responsible for an essential role in export or uptake of nocardichelin B (S2).

The rediscovery of the BGC for nocardichelin B showcases the potential of primary sequences and RetroMol. Retrieving and aligning biosynthetically similar compounds from NPAtlas for nocardichelin B succinctly highlights the uniqueness of its acyl starter as well as the overlap of its chelating repeats structure compared to other siderophores. Furthermore, fingerprinting of both the compound and target candidate clusters allowed for quick prioritization, using predicted modular building blocks, as well as the compound’s inferred siderophore identity.

## Discussion

Information about natural product biosynthesis and structure is scattered across heterogeneous data modalities and connected only by a relatively small number of well-studied natural products. RetroMol illustrates how multimodal integration and a joint embedding can address several central challenges in natural product discovery and characterization. By describing compounds and BGCs with a shared, biosynthetically informed language, RetroMol enables similarity-based retrieval among BGCs, natural product structures, and BGC-structure pairs, even when explicit links are missing.

RetroMol introduces novel functionality that should prove particularly useful given the rapidly expanding biosynthetic space revealed by large-scale genome sequencing. Systematic analyses have demonstrated that known natural products account for only a small fraction (3%) of the BGC diversity encoded in microbial genomes, and that biosynthetic novelty is unevenly distributed across taxonomic groups, with additional evidence for biome-specific diversity^36^. The wealth of uncharacterized BGCs underscores the importance of prioritization as a central bottleneck in natural product discovery^37–39^. A shared encoding provides a principled mechanism for such prioritization by allowing BGCs and compounds to be ranked by novelty and proximity to known biochemistry within a unified representation. Additionally, this shared representation provides a foundation for the development of modular platforms for chemical synthesis based on biosynthetic logic. For example, Namitharan et al. demonstrated that RetroMol-derived primary sequences can be translated into a programmable synthetic strategy, enabling the total synthesis of the natural product (+)-neopeltolide^40^.

At present, the primary limitation of RetroMol is the need to explicitly curate retrobiosynthetic rules. The default ruleset is currently focused on modular polyketides and NRPs, with partial coverage of related compound classes, including RiPPs. However, the software has been constructed in such a way that it facilitates customization by manually expanding this ruleset to cover additional biosynthetic pathways of interest. Consequently, the quality of cross-modal retrieval is inherently coupled to the quality of the biosynthetic hypotheses encoded in the parsing rules used for both compounds and BGCs. Finally, the current version of the RetroMol BGC parser used for the analyses in this work does not mine stereochemistry predictions from antiSMASH-mined BGCs, and therefore, stereochemistry-aware matching is reserved for unimodal compound retrieval.

Overall, RetroMol introduces an accessible framework for the deorphanization and annotation of likewise modular natural products and their putatively encoding BGCs. Even imperfect matches yield actionable hypotheses about BGC products, demonstrating the practical value of cross-modal biosynthetic fingerprints.

Multimodal embeddings for BGCs and compounds align closely with advances in mass spectrometry-based metabolomics. Molecular networking approaches, most notably integrated in the GNPS(2) environment, organize MS/MS data into molecular families and enable large-scale propagation of putative annotations across related compounds^41^. To provide further context to these molecular networks, MolNetEnhancer further integrates *in silico* structural annotations and automated chemical-class predictions to contextualize these molecular networks^42^. In parallel, methods like learned spectral similarity metrics and spectrum-to-fingerprint or spectrum-to-structure models, including MS2DeepScore, Spec2Vec, DreaMS, DiffMS, and related approaches, improve the correspondence between spectral and structural similarity^43–46^. These methods generate abstract, comparable representations from MS/MS data that are well-suited for integration into embedding-based retrieval frameworks.

Looking ahead, RetroMol is best viewed as a component within a broader multimodal knowledge graph ecosystem^47^, in which rule-based parsing, learned predictors, and experimental data coexist. Notably, RetroMol uses a user-defined ruleset making the tool expandable to cover yet unknown modular natural products, as well as hybrid compounds containing both modular and non-modular biosynthetic components such as nocardichelin B. As compounds are linked to tandem mass spectrometry spectra via molecular networking and spectrum-based learning approaches, these spectra can, in turn, be connected to BGCs via compound-level embeddings. Additionally, any insights obtained from comparing compounds and BGCs biosynthetically can inspire novel ways to engineer BGCs. In this way, improvements in retrobiosynthetic rules and biosynthetic annotation can propagate across modalities, strengthening genome-metabolite-spectrum links and enabling more comprehensive navigation of biosynthetic space.

## Methods

RetroMol was written in Python 3.10 using RDKit 2025.9.1^48^ as its cheminformatics engine.

### Data collection

We compiled reference chemical and biosynthetic data from two primary resources. 36,454 natural product compound structures were retrieved from NPAtlas version 3.0^9^. 513 experimentally characterized biosynthetic gene clusters (BGCs) with at least two NRPS and/or type I PKS modules, including associated metadata and at least one link to a reported chemical product, were collected from MIBiG version 4.0^25^.

### Generating structural fingerprints for compounds

Structural fingerprints for compounds were created using RDKit’s Morgan fingerprint implementation^48^. Fingerprints were generated with default settings, radius 2, 2048 bits, and without considering stereochemistry, unless explicitly stated otherwise.

### Parsing compounds

#### Reaction and matching ruleset

RetroMol uses two complementary rulesets to build a reaction graph: a directed heterogeneous graph where reactant nodes link to product nodes through reaction nodes. RetroMol uses two rule types in a RuleSet to create this reaction graph: reaction rules and matching rules (S4-5). Reaction rules are defined by SMILES arbitrary target specification (SMARTS) patterns (ReactionRule) and generate retrosynthetic products during graph expansion. Matching rules are defined by Simplified Molecular Input Line Entry System (SMILES) patterns (MatchingRule) and are used to identify product nodes and decide whether further expansion should stop.

Matching rules are by default “terminal”, which means that retrosynthesis stops when a product node is identified with one of the matching rules. The user can set the matching rule to be terminal=False, allowing the node to be identified as well as further retrosynthesized.

Additionally, matching rules can be assigned a stereochemistry flag, which determines if the matching rule is only used when stereochemistry matching is enabled (with flag -c on the command line or by setting match_stereochemistry=True). Finally, users can set additional pseudonyms for a matching rule, which are used during biosynthetic fingerprinting. For example, the amino acids serine and glycine can both be given a more coarse pseudonym “amino acid”. Since both monomers have the same coarse “amino acid” token, they will force their biosynthetic fingerprints to be more similar.

Parsing a compound with reaction rules starts by creating a queue with only the input compound (i.e., the parent) enqueued. For each parent molecule, the parser first attempts to identify “uncontested” reaction sites (non-overlapping reactive atom sets) among rules flagged allowed_in_bulk; these are applied together in a single bulk step to produce products, while enforcing atom-tag masking so only the matched atoms are modified. If no uncontested bulk application succeeds, the parser falls back to contested expansion and exhaustively applies each reaction rule individually to the parent in the order of which the rules were supplied. Each application yields sanitized product tuples. Stereo-aware duplicate product sets are removed before enqueueing. A quick check is applied to see if any multi-fragment products were produced, which will throw an error and prompts the user to update their reaction rules to prevent this from happening.

At each node, the parser calls MolNode.identify, which iterates over matching rules in order and assigns the first identity whose pattern matches the molecule. A full-molecule match is enforced by a substructure match plus equal atom and bond counts. Stereochemistry can be required if match_stereochemistry is enabled. If a node is identified and the matched rule is marked terminal=True, expansion is halted at that node. Otherwise, the node continues to be expanded by reaction rules. Rule ordering, therefore, matters for identity assignment, while reaction rules drive the generation of new nodes and edges in the reaction graph.

The default RuleSet loaded by RuleSet.load_default() contains 94 reaction rules from rxn.yml and 412 matching rules from mxn.yml. When stereochemistry matching is enabled, an additional 76 polyketide monomer stereoisomers are included in the matching ruleset. The chiral polyketide set identifies polyketide units with distinct chiral centers.

#### Retrosynthetic parsing

RetroMol parses compounds either as single SMILES strings or in batch from SDF, CSV/TSV, or JSON/JSONL files. In batch mode, inputs are streamed to keep memory usage low: SDF records are read with RDKit’s SDMolSupplier^48^, tables are streamed in chunks via pandas.read_csv(…, chunksize=…), and JSON/JSONL is parsed incrementally (JSONL line-by-line; JSON arrays via ijson). Each record is converted to a (smiles, props) tuple, where props preserves all original fields/columns. By default, the SMILES column is named “smiles”; additional fields are retained in the output as properties, ensuring traceability from the input to the result.

Each SMILES is standardized in a Submission object before parsing. The pipeline replaces “[N]” with “N” to avoid RDKit parsing issues, constructs an RDKit Mol from SMILES, keeps only the largest fragment, neutralizes charges by default, optionally canonicalizes tautomers (default off), sanitizes the molecule, and retains stereochemistry by default. An InChIKey is computed, and each atom is tagged with a unique isotope label so atom identity can be tracked through reactions. Retrosynthetic parsing then builds a reaction graph by breadth-first expansion: reaction rules are applied to parent molecules, products are sanitized and stereochemistry reassigned, multi-fragment products are rejected, and products with overlapping atom tags are filtered out. Expansion stops at “terminal” identities defined by the matching rules, and newly discovered products are enqueued using a canonical, tagged SMILES hash to prevent duplicates. Stereo-aware matching of rules can be enabled with the -c flag on the command line or by setting match_stereochemistry=True. Fast SDF parsing (with flag --rdkit-fast on the command line) skips initial sanitization and removes hydrogens, with sanitization performed only when needed. Example reaction graphs are included in S3.

#### Calculating coverage metric for retrosynthesis results

Coverage is computed from atom-isotope tags assigned during submission of a compound to the RetroMol parsing pipeline. RetroMol collects the union of isotope tags from all identified molecule nodes in the reaction graph and compares these to the isotope tags present in the original root molecule. The coverage metric is the fraction of root tags recovered within identified nodes.

If the root molecule has no tags (e.g., an empty or invalid molecule), coverage is defined as 0.0. This metric, therefore, measures how much of the original molecule’s tagged atom set has been assigned to identified monomer nodes in the reaction graph.

#### Calculating linear readouts from retrosynthesis results

The synthesis graph used for the linear readout is extracted from the heterogeneous reaction graph (nodes can be either a reaction/product or a reaction) via extract_min_edge_synthesis_graph. The reaction graph is treated as an AND/OR graph where molecule nodes are OR nodes (choose at most one outgoing reaction edge) and reaction edges are AND nodes (all precursor molecules must be solved). Costs are computed by dynamic programming with memoization and a cycle guard: identified terminal molecules have zero cost, identified non-terminal leaves may stop with a penalty (set to nonterminal_leaf_penalty=100 in run_retromol by default to prevent nonterminal leafs), and unexpanded, unidentified leaves incur an unsolved_leaf_penalty=5.0 by default. Each outgoing reaction edge has a cost of edge_base_cost=0.25, plus small tie-breakers that prefer uncontested over contested edges and slightly penalize high-branching edges. The minimum-cost choice at each node is selected, and a subgraph is assembled that includes the root molecule, the chosen outgoing edge (if any), and all required precursor branches. The synthesis extraction is considered “solved” if the total cost is finite and below the unsolved-leaf penalty threshold.

The linear readout is then computed from the synthesis subgraph with LinearReadout.from_reaction_graph(root_enc, r.graph), where root_enc is the hash of the canonical isomeric tagged SMILES of the submission’s root molecule. The readout first constructs an assembly graph by mapping leaf monomer tag sets back to the root molecule’s atom-isotope tags and adding edges whenever a root bond connects atoms from different monomers. Each edge stores the underlying root-bond metadata (atom indices, tags, element symbols, and bond type/order). The assembly graph includes an explicit unassigned node for regions of the root that are not covered by any monomer, and overlapping tag assignments are disallowed. Example assembly graphs are included in S3.

For the linear paths, the assembly graph is filtered to keep only root bonds between C-C and C-N atom pairs and to retain isolated nodes. Connected components are extracted, and the longest monomer path is computed per component. The final LinearReadout object stores the assembly graph and the list of longest-path monomer sequences.

#### Parsing NPAtlas compounds

We applied RetroMol to 36,454 microbial natural products from NPAtlas version 3.0^9^ (Figure 2a); we note that 252 compounds failed to parse or timed out after 30 seconds. Chemical superclass annotations for the parsed compounds were generated by submitting every SMILES to the NPClassifier API (accessed May 19, 2026)^2^. When a compound had multiple superclass annotations, it was counted in each corresponding group.

### Parsing BGCs

RetroMol loads antiSMASH^7,8^ GenBank files via load_regions, which dispatches to parse_antismash_gbk. The dispatcher can be expanded to include parsers for GenBank files from different sources, although for now only an antiSMASH GenBank parser is included.

GenBank files are parsed with Biopython version 1.86^49^. RetroMol parses each region GenBank file into a list of its antiSMASH-defined candidate cluster Region objects with accompanying metadata: candidate cluster bounds, strand, qualifiers (by default only product type), CDS features, and aSDomain features. Each CDS feature is parsed into a Gene object, and every aSDomain feature is parsed into a Domain object and nested into its parent Gene object.

#### Forward synthetic parsing and calculating linear readouts

A linear readout is constructed per candidate cluster Region by linear_readout, which builds NRPS modules gene-wise and PKS modules region-wise in the order they are found in.

NRPS modules are assembled per gene using domains in biosynthetic order (reverse on reverse-strand genes). Each A domain anchors a module. The module window extends one domain left if the immediately preceding domain is a condensation domain (C domain), and extends right until the next A domain. The module’s bounds are the min/max domain coordinates in the window, and its anatomy records presence of C/T/E/MT/Ox/R/TE domains. If the A domain has substrate predictions in its annotations, the highest-score prediction is stored as tuple (name, SMILES, score).

PKS modules are assembled across the region using a biosynthetic domain stream in the order they are found in. Each KS domain anchors a module; the window runs to the next KS (exclusive) and is truncated if domains are >20kb from the KS end. The module’s domain types are classified for presence of AT and active KR/DH/ER domains (activity inferred from qualifiers like “inactive/truncated”). AT loading mode is set to cis if an AT exists in the window, trans if an upstream AT-only gene exists, else unknown. If cis and only DHt (DH domain variant predominantly associated with trans-AT PKS clusters^50^) is present, DH is treated as inactive. Module bounds are from min/max domain coordinates and module indices are tracked per gene.

#### Configured gene and domain models for mining modifiers

RetroMol exposes two extension points for inference: gene models and domain models. Users implement GeneInferenceModel (operates on a Gene) or DomainInferenceModel (operates on a Domain) by defining “predict(…) -> list[InferenceResult]”, assigning a name, and optionally using the built-in result(…) helper to emit labels, scores, and metadata. Models are registered via register_gene_model or register_domain_model, and annotate_region runs all registered models over each gene and each domain, storing outputs in gene.annotations or domain.annotations for downstream use.

For all analyses in this manuscript, one gene model and one domain model were created and registered for inference: a Pfam gene model based on Pfam clan CL0113 to detect glycosyltransferases and a PARAS domain model of inferring A domain substrate specificities in NRP BGCs^19,51^.

#### Pfam gene annotation model

RetroMol treats Pfam HMMs as gene-level inference models and uses them to annotate each gene’s protein sequence. Each HMM file is wrapped as a PfamModel with a label derived from the filename stem. For each gene, the model runs HMMER (pyhmmer.hmmscan^52^) against that single HMM with an E-value cutoff of 1e-5 and Pfam “gathering” cutoffs enabled. If a hit is found, the model emits one InferenceResult with target=GENE, label=“hmm label”, and a confidence score). These gene-level results are stored in gene.annotations, and the linear-readout step converts their labels into “modifiers” by appending all gene annotation labels to the readout’s modifiers list.

#### PARAS A domain inference model

PARAS^19^ is a domain-level model applied only to antiSMASH domains whose type is “AMP-binding” (i.e., the NRPS A domain). For each such domain, RetroMol uses PARAS to predict the domain’s substrate specificity using the all_substrates_model (https://zenodo.org/records/17224548). The RetroMol wrapper returns the top three substrates above a probability threshold of 0.1. These predictions, including their SMILES, are stored in the Domain annotations and used later when building NRPS modules, where the highest-confidence A domain prediction becomes the module’s monomer. The current RetroMol around PARAS removes stereochemistry from the predicted monomers as no other matching rules for monomers, apart from the polyketide ones, currently have stereoisomers included in the default rule set.

#### Selecting MIBiG BGC-compound pairs for retrieval

We selected 513 NRP and type I polyketide BGCs from MIBiG version 4.0^25^ (as classified by antiSMASH as either NRPS, T1PKS, or both) with at least one assigned product structure and at least two modular domains (i.e., an NRP domain with C, A, and PCP domains, or a polyketide domain with KS and cis/trans AT domains). When a BGC had multiple associated products, we selected the compound with the highest molecular weight to be the representative. When a BGC contained multiple candidate clusters, we selected the longest candidate clusters to be the representative.

### Calculating biosynthetic fingerprints

Biosynthetic fingerprints were calculated from RetroMol linear readouts by representing each monomer as a set of biosynthetic tokens. For compound-derived entries, each identified monomer contributed its matched rule name and associated pseudonyms. For BGC-derived entries, each NRP module contributed its predicted monomer and associated pseudonyms and each polyketide module contributed its β-carbon oxidation state and associated pseudonyms (Figure 1b). Names and pseudonyms for monomers found in either compounds or BGCs were harmonized to allow for vocabulary overlap between biosynthetic fingerprints from compounds and BGCs. All names and pseudonyms can be found in the default matching rules (S5).

Unidentified monomers were encoded as “UNK”. A vocabulary was constructed from the default RetroMol matching rule set, using the rule names and pseudonyms, and each primary sequence was encoded with the Fingerprinter as a 1024-dimensional float32 vector using two deterministic hashes per token. Token contributions were normalized within each monomer before accumulation, producing count-based biosynthetic fingerprints suitable for cosine similarity comparison.

### Alignment of primary sequences

Primary sequences were aligned as ordered monomer-token sequences. Pairwise alignment was performed with Biopython’s (version 1.86^49^) PairwiseAligner in global mode using a custom substitution matrix. Matrix scores were derived from Tanimoto similarities between Morgan fingerprints of RetroMol rule SMILES strings, calculated with radius 2 and 2048 bits without stereochemistry. Additional self-scores were assigned for “UNK” and coarse PK tokens, and predefined PK subtype matches were hard-coded. Internal gap opening and extension penalties were set to -0.5 and -0.1, respectively, while terminal gap penalties were reduced to -0.2 and -0.05.

For retrieval reranking, each database sequence was aligned to the query in both forward and reverse orientations, and the higher alignment score was retained.

#### Calculating normalized alignment scores

Normalized alignment scores were calculated by dividing the pairwise alignment score between a query sequence and a target sequence by the maximum possible self-alignment score of the query sequence.

### Database construction and retrieval

The retrieval database was implemented in DuckDB^53^ version 1.5.3 in Python with one record per RetroMol linear readout. Each entry stored a stable SHA-256 identifier, compound or BGC type, name, URL, raw source string, primary monomer sequence, and the corresponding 1024-dimensional biosynthetic fingerprint. NPAtlas-derived compound entries were populated from RetroMol JSONL output by parsing each linear readout path, calculating its fingerprint, and inserting it as a database entry. MIBiG-derived BGC entries were populated from accession-matched MIBiG JSON and GenBank files.

Retrieval was performed by querying the DuckDB table with a query fingerprint and ranking entries by DuckDB’s array_cosine_similarity, filtering by entry type. Cross-modal retrieval was performed by querying the database with a BGC fingerprint and calculating fingerprint similarities with only compound fingerprints. The 1000 closest matches, with their metadata including their primary sequences, were retrieved based on cosine similarity. These 1000 closest matches were consecutively reranked based on the pairwise alignment score of their primary sequence with the query primary sequence. Top-N performance was determined by calculating the maximum Tanimoto coefficient between the SMILES of the top-N reranked matches and the query-associated SMILES.

### Dimensionality reduction of biosynthetic fingerprints

Biosynthetic fingerprints were converted to a feature matrix and projected to two dimensions using UMAP (umap-learn version 0.5.3^54^) using cosine as metric, with n_neighbors and min_dist set to 15 and 0.5 respectively. The reduced dimensions were used as coordinates to generate the scatter plot in Figure 4c, with one point per biosynthetic fingerprint.

### Metabolomics data collection for nocardichelin B

Untargeted metabolomics data were generated for four *Nocardia* strains from the DTU New Bioactive Compounds (NBC) Collection: *Nocardia* sp. NBC_01329 (CP108381), NBC_00511 (CP107850), NBC_01009 (CP108695), and NBC_01730 (CP109162)^31^. For the latter three, all were prepared in liquid culture, whereas NBC_01329 was grown and extracted from solid cultures.

For NBC_00511, NBC_01009, and NBC_01730, precultures were grown in ISP2 (4 g L⁻¹ yeast extract (Bacto, BD Biosciences, USA), 4 g L⁻¹ dextrose, 10 g L⁻¹ malt extract (Sigma-Aldrich)) and inoculated 30 μL into four 100 mL Erlenmeyer flasks containing 30 mL of MA (beef extract replaced with meat extract)^55^, ISP2, DNPM^56^, and SoyM (10 g L⁻¹ mannitol, 10 g L⁻¹ soy flour (W. Schoenenberger GmbH, Magstadt, Germany); half the standard concentration). Cultures were incubated at 30 °C and 140 rpm, with a metal spring to improve aeration. NBC 00511 and NBC_01730 were grown for 4 days, NBC_01009 for 2 days. Supernatants (2mL) were collected after centrifugation at 5000 × g (5 min) and stored at -20 °C.

NBC_01329 was cultured on solid media (MA, ISP2, DNPM, SoyM with 20 g L⁻¹ mannitol and soy flour, plus 18 g L⁻¹ agar). After autoclaving, 30 mL of each medium was poured into 90 mm petri dishes. Once solidified, 7 mm agar plugs were punched, and three plugs per medium were transferred to a 12-well plate. A preculture (ISP2) was pelleted, washed, and resuspended in sterile water; 10 μL was inoculated onto each plug. Plates were incubated at 30 °C for 4 days. Agar plugs were then extracted twice with 400 µL of sterile water (10 minutes each) and twice with methanol. Methanol extracts were evaporated and combined with water extracts prior to being stored at -20 °C.

All samples were then processed using an Oasis HLB 96-well plate (30 mg sorbent; Waters) using a Waters Positive Pressure-96 Processor system. The protocol included conditioning (1 mL methanol, 1 mL water), sample loading (600 μL), washing (600 μL 15% methanol), and elution (600 μL 40%, 70%, and 100% methanol). The 70% and 100% fractions were dried in a vacuum centrifuge and reconstituted in 75 μL 50% methanol.

LC-MS/MS analyses were performed on either a Vanquish Duo UHPLC system coupled to an Orbitrap ID–X mass spectrometer or a Dionex Ultimate 3000 UHPLC system coupled to a Fusion MS (Thermo Fisher Scientific, USA) equipped with a heated electrospray ionisation (HESI). Chromatographic separation was achieved on a Waters ACQUITY BEH C18 column (2.1 × 100 mm, 1.7 μm) (WatersTM, Milford, MA, USA) maintained at 40 °C. Mobile phase A was MilliQ water containing 0.1% formic acid (VWR Chemicals, USA), and mobile phase B was LC-MS grade acetonitrile (VWR Chemicals) containing 0.1% formic acid delivered at 0.35 mL/min. The gradient started at 2% B (0.8 min), increased linearly to 5% B (3.1 min), then to 100% B (6.7 min), held for 1 min, and equilibrated for 2.7 min. The injection volume was 1 µL.

Mass spectrometry data were acquired in positive HESI mode (3.5 kV) over m/z 100-1500 and in profile mode. Full MS and data-dependent MS/MS scans were collected at resolutions of 120,000 and 30,000, respectively. The top ten precursor ions per MS scan were isolated (1.6 m/z window) and fragmented by higher-energy collisional dissociation (HCD) using stepped normalized collision energies of 20, 40, and 60%. Dynamic exclusion was set at 6 s (±6 ppm). The automatic gain control (AGC) targets were set to 4 × 10⁵ for full MS and 5 × 10⁴ for MS/MS.

### Metabolomics and BGC data analysis for nocardichelin B

Raw data files were converted to mzML format using MSConvert (ProteoWizard version 3.0) and uploaded to the GNPS2 platform (https://gnps2.org/) for molecular networking analysis^41^. Classical Molecular Networking was performed using GNPS2 (release 2025.08.06) and modifications between spectra were analyzed with ModiFinder^57^. The minimum cluster size was set to 1, and precursor ion and fragment ion mass tolerances were both set to 0.01 Da. Molecular networks were visualised and annotated using Cytoscape version 3.10^58^. Supporting files for the metabolomics analysis are provided in S2, and the data were deposited in MASSIVE (MSV000100273).

Whole-genome sequences of the four *Nocardia* strains were analysed using antiSMASH v7.0 with default parameters^7^. All region GenBank files for the identified BGCs from the four *Nocardia* strains were parsed with RetroMol, along with the SMILES for Nocardichelin B from NPAtlas (NPA007982)^9^. Candidate cluster two from the uploaded GenBank file for region twenty-two (basepairs 4,889,794 to 5,001,770) in *Nocardia sp.* Strain NBC_01329 (the second nearest neighbor to the nocardichelin B biosynthetic fingerprint readout) provided the best match upon manual inspection of the ten nearest candidate clusters in fingerprint space. The candidate BGC from strain NBC_01329 was compared to the recently published Nocardichlin B BGC using clinker v0.0.28^59^.

## Supporting information

RetroMol Supplemental Information

## Data availability

All the generated LC-MS files used for the rediscovery of the nocardichelin B BGC have been deposited into MassIVE (dataset identifier MSV000100273).

RetroMol is available on GitHub: https://github.com/moltools/retromol.

## References

1. Medema, M. H., De Rond, T. & Moore, B. S. Mining genomes to illuminate the specialized chemistry of life. Nat. Rev. Genet. 22, 553–571 (2021).

2. Kim, H. W. et al. NPClassifier: A Deep Neural Network-Based Structural Classification Tool for Natural Products. J. Nat. Prod. 84, 2795–2807 (2021).

3. Fischbach, M. A. & Walsh, C. T. Assembly-Line Enzymology for Polyketide and Nonribosomal Peptide Antibiotics: Logic, Machinery, and Mechanisms. Chem. Rev. 106, 3468–3496 (2006).

4. Terlouw, B. R. et al. RAIChU: automating the visualisation of natural product biosynthesis. J. Cheminformatics 16, 106 (2024).

5. Rutz, A. et al. MITE: the Minimum Information about a Tailoring Enzyme database for capturing specialized metabolite biosynthesis. Nucleic Acids Res. gkaf969 (2025) doi:10.1093/nar/gkaf969.

6. Butler, M. S. & Capon, R. J. Natural product inspired antibiotics approved for human use – 1943 to 2025. Nat. Prod. Rep. 10.1039.D5NP00067J (2026) doi:10.1039/D5NP00067J.

7. Blin, K. et al. antiSMASH 7.0: new and improved predictions for detection, regulation, chemical structures and visualisation. Nucleic Acids Res. 51, W46–W50 (2023).

8. Blin, K. et al. antiSMASH 8.0: extended gene cluster detection capabilities and analyses of chemistry, enzymology, and regulation. Nucleic Acids Res. 53, W32–W38 (2025).

9. Poynton, E. F. et al. The Natural Products Atlas 3.0: extending the database of microbially derived natural products. Nucleic Acids Res. 53, D691–D699 (2025).

10. Olkhovskii, I., Kushnareva, A., Popov, P., Tagirdzhanov, A. & Gurevich, A. Nerpa 2: linking biosynthetic gene clusters to nonribosomal peptide structures. Preprint at 10.1101/2024.11.19.624380 (2024).

11. Chainani, Y. et al. Merging the computational design of chimeric type I polyketide synthases with enzymatic pathways for chemical biosynthesis. Nat. Commun. 16, 5787 (2025).

12. Yan, D. et al. Seq2Saccharide: Discovering Oligosaccharides and Aminoglycosides Natural Products by Integrating Computational Mass Spectrometry and Genome Mining. J. Am. Chem. Soc. 147, 35323–35338 (2025).

13. Skinnider, M. A., Merwin, N. J., Johnston, C. W. & Magarvey, N. A. PRISM 3: expanded prediction of natural product chemical structures from microbial genomes. Nucleic Acids Res. 45, W49–W54 (2017).

14. Dejong, C. A. et al. Polyketide and nonribosomal peptide retro-biosynthesis and global gene cluster matching. Nat. Chem. Biol. 12, 1007–1014 (2016).

15. Larralde, M. & Zeller, G. Machine learning inference of natural product chemistry across biosynthetic gene cluster types. Preprint at 10.1101/2025.03.13.642868 (2025).

16. Khater, S., Anand, S. & Mohanty, D. In silico methods for linking genes and secondary metabolites: The way forward. Synth. Syst. Biotechnol. 1, 80–88 (2016).

17. Draisma, A., et al. BiG-SCAPE 2.0 and BiG-SLiCE 2.0: scalable, accurate and interactive sequence clustering of metabolic gene clusters.

18. Boldini, D. et al. Effectiveness of molecular fingerprints for exploring the chemical space of natural products. J. Cheminformatics 16, 35 (2024).

19. Terlouw, B. R. et al. PARAS: High-Accuracy Machine Learning of Substrate Specificities in Nonribosomal Peptide Synthetases. JACS Au 6, 2315–2336 (2026).

20. Healy, A. R., Vinale, F., Lorito, M. & Westwood, N. J. Total Synthesis and Biological Evaluation of the Tetramic Acid Based Natural Product Harzianic Acid and Its Stereoisomers. Org. Lett. 17, 692–695 (2015).

21. Nollen, L.-M., Meijer, D., Sorokina, M. & Van Der Hooft, J. Biosynfoni: A Biosynthesis-informed and Interpretable Lightweight Molecular Fingerprint. Preprint at 10.26434/chemrxiv-2025-cwq74 (2025).

22. Paterson, I., Britton, R., Delgado, O., Meyer, A. & Poullennec, K. G. Total Synthesis and Configurational Assignment of (−)-Dictyostatin, a Microtubule-Stabilizing Macrolide of Marine Sponge Origin. Angew. Chem. Int. Ed. 43, 4629–4633 (2004).

23. Canales, A. et al. The Bound Conformation of Microtubule-Stabilizing Agents: NMR Insights into the Bioactive 3D Structure of Discodermolide and Dictyostatin. Chem. – Eur. J. 14, 7557–7569 (2008).

24. Jungmann, K. et al. Two of a Kind—The Biosynthetic Pathways of Chlorotonil and Anthracimycin. ACS Chem. Biol. 10, 2480–2490 (2015).

25. Zdouc, M. M. et al. MIBiG 4.0: advancing biosynthetic gene cluster curation through global collaboration. Nucleic Acids Res. 53, D678–D690 (2025).

26. Butler, T. et al. MS2Mol: A transformer model for illuminating dark chemical space from mass spectra. Preprint at 10.26434/chemrxiv-2023-vsmpx-v3 (2023).

27. Jungmann, K. et al. Two of a Kind—The Biosynthetic Pathways of Chlorotonil and Anthracimycin. ACS Chem. Biol. 10, 2480–2490 (2015).

28. Chen, H., O’Connor, S., Cane, D. E. & Walsh, C. T. Epothilone biosynthesis: assembly of the methylthiazolylcarboxy starter unit on the EpoB subunit. Chem. Biol. 8, 899–912 (2001).

29. Medema, M. H. et al. Pep2Path: Automated Mass Spectrometry-Guided Genome Mining of Peptidic Natural Products. PLoS Comput. Biol. 10, e1003822 (2014).

30. Schneider, K. et al. Nocardichelins A and B, Siderophores from *Nocardia* Strain Acta 3026. J. Nat. Prod. 70, 932–935 (2007).

31. Jørgensen, T. S. et al. A treasure trove of 1034 actinomycete genomes. Nucleic Acids Res. 52, 7487–7503 (2024).

32. Fisk, M. B., Barrera Ramirez, J., Merrick, C. E., Wencewicz, T. A. & Gulick, A. M. Identification and Characterization of the Biosynthesis of the Hybrid NRPS-NIS Siderophore Nocardichelin. ACS Chem. Biol. 20, 1435–1446 (2025).

33. Shahneh, M. R. Z. et al. ModiFinder: Tandem Mass Spectral Alignment Enables Structural Modification Site Localization. J. Am. Soc. Mass Spectrom. 35, 2564–2578 (2024).

34. Maglangit, F. et al. Characterization of the promiscuous *N* -acyl CoA transferase, LgoC, in legonoxamine biosynthesis. Org. Biomol. Chem. 18, 2219–2222 (2020).

35. Butterton, J. R. & Calderwood, S. B. Identification, cloning, and sequencing of a gene required for ferric vibriobactin utilization by Vibrio cholerae. J. Bacteriol. 176, 5631–5638 (1994).

36. Gavriilidou, A. et al. Compendium of specialized metabolite biosynthetic diversity encoded in bacterial genomes. Nat. Microbiol. 7, 726–735 (2022).

37. Jo, W. S., et al. Total Synthesis and Structural Revision of Rhabdobranin Reveals a Cryptic Gram-Negative Antibiotic. Preprint at 10.26434/chemrxiv.15003454/v1 (2026).

38. Kalkreuter, E., Pan, G., Cepeda, A. J. & Shen, B. Targeting Bacterial Genomes for Natural Product Discovery. Trends Pharmacol. Sci. 41, 13–26 (2020).

39. Malit, J., Leung, H. & Qian, P.-Y. Targeted Large-Scale Genome Mining and Candidate Prioritization for Natural Product Discovery. Mar. Drugs 20, 398 (2022).

40. Namitharan, K., Saney, L., Filatov, V., Piano, F. & Healy, A. R. Synthesis of (+)-Neopeltolide via Sequence-Guided Iterative Assembly. Preprint at 10.26434/chemrxiv.15002531/v1 (2026).

41. Aron, A. T. et al. Reproducible molecular networking of untargeted mass spectrometry data using GNPS. Nat. Protoc. 15, 1954–1991 (2020).

42. Ernst, M. et al. MolNetEnhancer: Enhanced Molecular Networks by Integrating Metabolome Mining and Annotation Tools. Metabolites 9, 144 (2019).

43. Huber, F., Van Der Burg, S., Van Der Hooft, J. J. J. & Ridder, L. MS2DeepScore: a novel deep learning similarity measure to compare tandem mass spectra. J. Cheminformatics 13, 84 (2021).

44. Huber, F. et al. Spec2Vec: Improved mass spectral similarity scoring through learning of structural relationships. PLOS Comput. Biol. 17, e1008724 (2021).

45. Bushuiev, R. et al. Self-supervised learning of molecular representations from millions of tandem mass spectra using DreaMS. Nat. Biotechnol. 10.1038/s41587-025-02663-3 (2025) doi:10.1038/s41587-025-02663-3.

46. Bohde, M., Manjrekar, M., Wang, R., Ji, S. & Coley, C. W. DiffMS: Diffusion Generation of Molecules Conditioned on Mass Spectra. Preprint at 10.48550/arXiv.2502.09571 (2025).

47. Meijer, D. et al. Empowering natural product science with AI: leveraging multimodal data and knowledge graphs. Nat. Prod. Rep. 42, 654–662 (2025).

48. Landrum, G. et al. rdkit/rdkit: 2025_09_1 (Q3 2025) Release. Zenodo 10.5281/zenodo.17232453 (2025).

49. Cock, P. J. A. et al. Biopython: freely available Python tools for computational molecular biology and bioinformatics. Bioinformatics 25, 1422–1423 (2009).

50. NRPS/PKS domains - antiSMASH Documentation. https://docs.antismash.secondarymetabolites.org/modules/nrps_pks_domains/.

51. Liu, J. & Mushegian, A. Three monophyletic superfamilies account for the majority of the known glycosyltransferases.

52. Larralde, M. & Zeller, G. PyHMMER: a Python library binding to HMMER for efficient sequence analysis. Bioinformatics 39, btad214 (2023).

53. Raasveldt, M. & Mühleisen, H. DuckDB: an Embeddable Analytical Database. in Proceedings of the 2019 International Conference on Management of Data 1981–1984 (ACM, Amsterdam Netherlands, 2019). doi:10.1145/3299869.3320212.

54. McInnes, L., Healy, J., Saul, N. & Großberger, L. UMAP: Uniform Manifold Approximation and Projection. J. Open Source Softw. 3, 861 (2018).

55. Román-Hurtado, F. et al. One Pathway, Two Cyclic Non-Ribosomal Pentapeptides: Heterologous Expression of BE-18257 Antibiotics and Pentaminomycins from Streptomyces cacaoi CA-170360. Microorganisms 9, 135 (2021).

56. Jönsson, M. et al. Machine learning uncovers the transcriptional regulatory network for the production host Streptomyces albidoflavus. Cell Rep. 44, 115392 (2025).

57. Shahneh, M. R. Z. et al. ModiFinder: Tandem Mass Spectral Alignment Enables Structural Modification Site Localization. J. Am. Soc. Mass Spectrom. 35, 2564–2578 (2024).

58. Shannon, P. et al. Cytoscape: A Software Environment for Integrated Models of Biomolecular Interaction Networks. Genome Res. 13, 2498–2504 (2003).

59. Gilchrist, C. L. M. & Chooi, Y.-H. clinker & clustermap.js: automatic generation of gene cluster comparison figures. Bioinformatics 37, 2473–2475 (2021).

