## Supplementary material for "RetroMol: Parsing a shared encoding from natural products and their biosynthetic gene clusters": RetroMol Supplemental Information: S4_default_reaction_rules.yml

```
- name: break ester bond (intermolecular)
  smarts: "[*:6]-[C,c:1]-[O;!R:2]-[C;!R:3](=[O:4])-[C,c:5]>>[*:6]-[C,c:1]-[OH:2].[O]-[C:3](=[O:4])-[C,c:5]"
  allowed_in_bulk: true

- name: break thio-ester bond (intermolecular)
  smarts: "[*:6]-[C,c:1]-[S;!R:2]-[C;!R:3](=[O:4])-[C,c:5]>>[*:6]-[C,c:1]-[SH:2].[O]-[C:3](=[O:4])-[C,c:5]"
  allowed_in_bulk: true

- name: break N-O glycosidic bond (intermolecular)
  smarts: "[C:1][NH1:2][O:3][C:4]>>[C:1][NH2:2].[OH:3][C:4]"
  allowed_in_bulk: true

- name: break glycosidic bond
  smarts: "[C,c:1][O:2][C:3]1[O:4][C:5][C:6][C:7][C:8]1>>[C,c:1][OH:2].[OH][C:3]1[O:4][C:5][C:6][C:7][C:8]1"
  allowed_in_bulk: true

- name: break furanosidic bond
  smarts: "[C,c:1][O:2][C:3]1[O:4][C:5][C:7][C:8]1>>[C,c:1][OH:2].[OH][C:3]1[O:4][C:5][C:7][C:8]1"
  allowed_in_bulk: true

- name: break glycosidic ester bond
  smarts: "[C:8](=[O:9])[O:7][C:1]1[C:2][O:3][C:4][C:5][C:6]1>>[C:8](=[O:9])[OH:7].[O][C:1]1[C:2][O:3][C:4][C:5][C:6]1"
  allowed_in_bulk: true

- name: break N-glycosidic linkage
  smarts: "[O:2][C:3]1[O:4][C:5][C:6]([N:9][C:10])[C:7][C:8]1>>[O:2][C:3]1[O:4][C:5][C:6]([N:9])[C:7][C:8]1.[O][C:10]"
  allowed_in_bulk: true

- name: reverse epoxidation
  smarts: "[C:1]1[O:2][C:3]1>>[C:1]=[C:3].[O:2]"
  allowed_in_bulk: true

- name: reverse O-methylation
  smarts: "[O&D2:1][CH3:2]>>[O:1].[C:2]"
  allowed_in_bulk: true

- name: reverse N-methylation
  smarts: "[N:1][CH3:2]>>[N:1].[C:2]"
  allowed_in_bulk: true

- name: reverse S-methylation
  smarts: "[S:1][CH3:2]>>[S:1].[C:2]"
  allowed_in_bulk: true

- name: reverse C-methylation amino acid
  smarts: "[N:1][C:2]([CH3:3])([*:4])[C:5](=[O:6])[O:7]>>[N:1][C:2]([*:4])[C:5](=[O:6])[O:7].[CH3:3]"
  allowed_in_bulk: true

- name: reverse halogenation
  smarts: "[C,c:1]([Cl,Br,F,I:2])>>[C,c:1].[Cl,Br,F,I:2]"
  allowed_in_bulk: true

- name: reverse carbamic acid tailoring (type 1)
  smarts: "[NH2:1]-[CH0:2](=[O:3])-[OH0:4]-[*:5]>>[OH1:4]-[*:5].[NH2:1]-[CH0:2](=[O:3])[OH]"
  allowed_in_bulk: true

- name: reverse carbamic acid tailoring (type 2)
  smarts: "[NH2:1]-[C:2](=[O:3])-[N:4]-[*:5]>>[N:4]-[*:5].[NH2:1]-[C:2](=[O:3])[OH]"
  allowed_in_bulk: true

- name: reverse carbinolamide tailoring
  smarts: "[OH:1]-[C:2]1[C:3][C:4][O:5][C:6](=[O:7])[N:8]1>>[O:1]=[C:2][C:3][C:4][OH:5].[O][C:6](=[O:7])[N:8]"
  allowed_in_bulk: true
  props:
    references:
      - "PMID:12060743"

- name: reverse acetic acid tailoring
  smarts: "[CH3:1]-[CH0:2](=[O:3])-[OH0:4]-[*:5]>>[OH1:4]-[*:5].[CH3:1]-[CH0:2](=[O:3])[OH]"
  allowed_in_bulk: true

- name: reverse carboxylic acid tailoring
  smarts: "[OH1:1]-[CH0:2](=[O:3])-[NH1:4]-[*:5]>>[NH2:4]-[*:5].[OH1:1]-[CH0:2](=[O:3])[OH]"
  allowed_in_bulk: true

- name: reverse acylation (type 1)
  smarts: "[CH3:1][C:2](=[O:3])[O,N:4][*:5]>>[CH3:1][C:2](=[O:3])[OH].[O,N:4][*:5]"
  allowed_in_bulk: true

- name: reverse acylation (type 2)
  smarts: "[CH1:1]([CH3:6])([CH3:7])[C:2](=[O:3])[O,N:4][*:5]>>[CH1:1]([CH3:6])([CH3:7])[C:2](=[O:3])[OH].[O,N:4][*:5]"
  allowed_in_bulk: true

- name: reverse acylation (type 3)
  smarts: "[*:1][NH1:2][CH2:3][CH3:4]>>[*:1][NH2:2].[OH][CH2:3][CH3:4]"
  allowed_in_bulk: true

- name: reverse cyanide tailoring
  smarts: "[C:1][C&D2:2]#[N&D1:3]>>[C:1].[C:2]#[N:3]"
  allowed_in_bulk: true

- name: reverse hydroxyl sulfonation
  smarts: "[*:1][S:2](=[O:3])(=[O:4])[OH:5]>>[*:1].[OH][S:2](=[O:3])(=[O:4])[OH:5]"
  allowed_in_bulk: true

- name: reverse hydroxyl phosphonylation
  smarts: "[C:3][O:1][P:2]>>[C:3][O:1].[O][P:2]"
  allowed_in_bulk: true

- name: reverse 1,3-dioxane formation (cyclic acetal)
  smarts: "[*:6][C:1]1~[C:2]~[C:3]([*:7])[O:4][C:8][O:5]1>>[*:6][C:1]([OH:5])~[C:2]~[C:3]([OH:4])[*:7].[C:8]=[O]"

- name: reverse 1,3-dioxolane formation (spiro-bis(pyranose) system)
  smarts: "[*:6][C:1]1~[C:2]([*:7])[O:4][C:8][O:5]1>>[*:6][C:1]([OH:5])~[C:2]([OH:4])[*:7].[C:8][OH]"

- name: reverse 2,3-diaminopropionate amidation
  smarts: "[N:1][CH1:2]([C:3](=[O:4])[NH2:5])[CH2:6][NH2:7]>>[NH1:1][CH1:2]([C:3](=[O:4])[OH])[CH2:6][NH2:7].[N:5]"

- name: reverse hydrogenation tailoring
  smarts: "[CH2:1][CH1:2]([OH1:3])[CH2:4][OH1:5]>>[CH2:1][CH0:2](=[OH0:3])[OH1].[CH3:4][OH1:5]"

- name: reverse etherification
  smarts: "[*:1]-[CH0:2]1(-[OH:3])-[O:4]-[CH1:5](-[*:6])-[C:7][C:8][C:9]1>>([*:1]-[CH0:2]1=[OH0:3].[OH1:4]-[CH1:5](-[*:6])-[C:7][C:8][C:9]1)"

- name: reverse terminal etherification
  smarts: "[CH1:2]1(-[OH:3])-[O:4]-[CH1:5](-[*:6])-[C:7][C:8][C:9]1>>([CH0:2]1(=[OH0:3])[OH].[OH1:4]-[CH1:5](-[*:6])-[C:7][C:8][C:9]1)"

- name: reverse etherification (6-ring; no hydroxyl group)
  smarts: "[C:1]-[C:2]([C:5]~[C:6]~[C:7]1)-[O:3]-[C:4]1[C:8][C:9]>>([C:1]-[C:2]([C:5][C:6][C:7]1)-[OH0:3].[C:4]1=[C:8][C:9])"

- name: reverse etherification (5-ring; no hydroxyl group)
  smarts: "[C:1]-[C:2]([C:5]~[C:7]1)-[O:3]-[C:4]1[C:8][C:9]>>([C:1]-[C:2]([C:5][C:7]1)-[OH0:3].[C:4]1=[C:8][C:9])"

- name: reverse macrolactonization
  smarts: "[C;R:1][C;R:2](=[O:3])[O;R:4][C;R:5]>>([C:1][C:2](=[O:3])[OH].[OH:4][C:5])"

- name: break ester bond (intermolecular)
  smarts: "[C,c:1][C;!R:2](=[O:3])[O;!R:4][C:5]>>[C,c:1][C:2](=[O:3])[OH].[OH:4][C:5]"

- name: break aromatic ester bond (intramolecular)
  smarts: "[c:1]1[c:2][c:3][c:4][c:5](=[O:7])[o:6]1>>([OH:6][C:1]=[C:2]-[C:3]=[C:4]-[C:5](=[O:7])[OH])"

- name: reverse macrolactonethionization
  smarts: "[C;R:1][C;R:2](=[O:3])[S;R:4][C;R:5]>>([C:1][C:2](=[O:3])[OH].[SH:4][C:5])"

- name: break thio-ester bond (intermolecular)
  smarts: "[C:1][C;!R:2](=[O:3])[S;!R:4][C:5]>>[C:1][C:2](=[O:3])[OH].[SH:4][C:5]"

- name: reverse carbocyclization (type 1)
  smarts: "[*:1][C:2]1[C:3]([*:4])[C:5]=[C:6][CH1:7]([C;R:8])[C:9]([C;R:10])1>>([*:1][C:2]=1.[C:3]([*:4])=[C:5]-[C:6]=[CH1:7]([C;R:8]).[C:9]([C;R:10])=1)"

- name: reverse carbocyclization (type 2)
  smarts: "[*:1][C:2]1[C:3]([*:4])[C:5]=[C:6][C:7](=[C;R:8][C;R:11])[C:9]([C;R:10])1>>([*:1][C:2]=1.[C:3]([*:4])=[C:5]-[C:6]=[C:7](-[C;R:8]=[C;R:11]).[C:9]([C;R:10])=1)"

- name: reverse carbocycle oxidation
  smarts: "[#6:1]1(-[#6:4](=[#6:6]-[#6:7](-[#8:3])-[#6:8](-[#6:10])-[#6:2]-1-[#6:9])-[*:11])-[*:12]>>[#6:1]1(-[#6:4](-[#6:6]=[#6:7]-[#6:8](-[#6:10])-[#6:2]-1-[#6:9])-[*:11])-[*:12].[#8:3]"

- name: reverse heterocyclization
  smarts: "[C:1]1[C:2][N:3][C:4](=[O:5])[C:6]=1([*:7])>>[C:1]([OH])(=[O])[C:2][N:3][C:4](=[O:5])[C:6]([*:7])"

- name: break dilsufide bridge (intramolecular)
  smarts: "[C;R:1][C:2][S:3][S:4][C:5][C;R:6]>>([C;R:1][C:2][SH:3].[SH:4][C:5][C;R:6])"
  allowed_in_bulk: true

- name: break disulfide bridge (intermolecular)
  smarts: "[C:1][C:2][S:3][S:4][C:5][C:6]>>[C;R:1][C:2][SH:3].[SH:4][C:5][C;R:6]"
  allowed_in_bulk: true

- name: break threonine-cysteine bridge (intramolecular)
  smarts: "[C:1](=[O:2])[C:3]([N:4])[CH1:5]([CH3:6])[S:7][CH2:8][C:9]([N:10])[C:11]=[O:12]>>([C:1](=[O:2])[C:3]([N:4])[CH1:5]([CH3:6])[OH].[SH:7][CH2:8][C:9]([N:10])[C:11]=[O:12])"
  allowed_in_bulk: true

- name: break threonine-cysteine bridge (intermolecular)
  smarts: "[C:1](=[O:2])[C:3]([N:4])[CH1:5]([CH3:6])[S:7][CH2:8][C:9]([N:10])[C:11]=[O:12]>>[C:1](=[O:2])[C:3]([N:4])[CH1:5]([CH3:6])[OH].[SH:7][CH2:8][C:9]([N:10])[C:11]=[O:12]"
  allowed_in_bulk: true

- name: break serine-cysteine bridge (intramolecular)
  smarts: "[C:1](=[O:2])[C:3]([N:4])[CH2:5][S:7][CH2:8][C:9]([N:10])[C:11]=[O:12]>>([C:1](=[O:2])[C:3]([N:4])[CH2:5][SH:7].[OH][CH2:8][C:9]([N:10])[C:11]=[O:12])"
  allowed_in_bulk: true

- name: break serine-cysteine bridge (intermolecular)
  smarts: "[C:1](=[O:2])[C:3]([N:4])[CH2:5][S:7][CH2:8][C:9]([N:10])[C:11]=[O:12]>>[C:1](=[O:2])[C:3]([N:4])[CH2:5][SH:7].[OH][CH2:8][C:9]([N:10])[C:11]=[O:12]"
  allowed_in_bulk: true

- name: break cysteine bond (intramolecular)
  smarts: "[O:1][C:2](=[O:3])[C:4][C:5][SH0:6][C:7]>>[O:1][C:2](=[O:3])[C:4][C:5][SH1:6].[C:7]"
  allowed_in_bulk: true

- name: reverse reduction (type 1)
  smarts: "[NH2:1][CH0:2]([OH:3])[C:4](=[O:5])[OH:6]>>[NH2:1][CH1:2][C:4](=[O:5])[OH:6].[OH2:3]"

- name: reverse reduction (type 2)
  smarts: "[NH2:1][CH0:2]([OH:3])[CH2:4][C:5](=[O:6])[OH:7]>>[NH2:1][CH1:2][CH2:4][C:5](=[O:6])[OH:7].[OH2:3]"

- name: reverse oxazole formation
  smarts: "[C,c:1][c:2]1[o:3][c:4][c:5]([C:6])[n:7]1>>[C,c:1][CH0:2](=[O:3])[NH1:7][CH1:5]([CH2:4][OH1])[C:6]"

- name: reverse oxazoline formation
  smarts: "[C,c:1][C:2]=1[OH0:3][C:4][C:5]([C:6])[N:7]1>>[C,c:1][CH0:2](=[O:3])[NH1:7][CH1:5]([CH2:4][OH1])[C:6]"

- name: reverse thiazole formation
  smarts: "[C:1][c:2]1[s:3][c:4][c:5][n:7]1>>([C:1]-[CH0:2]-1(=O).[SH1:3]-[CH2:4]-[CH1:5]-[NH1:7]-1)"

- name: reverse thiazoline tautomerization
  smarts: "[OH0:1]=[C:2]1-[S:3]-[C:4]-[C:5](-[C:6])-[N:7]-1>>[OH1:1]-[C:2]1-[S:3]-[C:4]-[C:5](-[C:6])-[N:7]=1"

- name: reverse thiazoline formation (type 1)
  smarts: "[*:1][C:2]=1[SH0:3][C:4][C:5][N:7]1>>([*:1]-[CH0:2]-1(=O).[SH1:3]-[CH2:4]-[C:5]-[NH1:7]-1)"

- name: reverse thiazoline formation (type 2)
  smarts: "[*:1][C:2]([C:3][SH0:4]1)[NH1:5][C:6]1[*:7]>>[*:1][C:2]([C:3][SH1:4])[NH1:5][CH0:6](=[O])[*:7]"

- name: reverse tetramate formation (type 1)
  smarts: "[*:1]-[C:2](-[OH:3])=[C:4]1[C:5](=[O:6])[N:7][C:8][C:9]1=[O:10]>>[*:1]-[C:2](=[OH0:3])-[CH2:4][C:5](=[O:6])[N:7][C:8][C:9](-[OH])=[O:10]"

- name: reverse tetramate formation (type 2)
  smarts: "[*:1]-[C:2](-[OH:3])-[C:4]=1[C:5](=[O:6])[N:7][C:8][C:9]1-[OH:10]>>[*:1]-[C:2](=[OH0:3])-[CH2:4][C:5](=[O:6])[N:7][C:8][C:9](-[OH])=[O:10]"

- name: reverse tetramate formation (type 3; aromatic)
  smarts: "[*:1]-[C:2](=[O:3])[c:4]1[c:5](-[OH:6])[n:7][c:8][c:9]1-[OH:10]>>[*:1]-[C:2](=[OH0:3])-[CH2:4][C:5](=[O:6])[N:7][C:8][C:9](-[OH])=[O:10]"

- name: reverse tetronate formation (type 1)
  smarts: "[*:1]-[C:2](-[OH:3])=[C:4]1[C:5](=[O:6])[O:7][C:8][C:9]1=[O:10]>>[*:1]-[C:2](=[OH0:3])-[CH2:4][C:5](=[O:6])[O:7][C:8][C:9](-[OH])=[O:10]"

- name: reverse tetronate formation (type 2)
  smarts: "[*:1]-[C:2](-[OH:3])-[C:4]=1[C:5](=[O:6])[O:7][C:8][C:9]1-[OH:10]>>[*:1]-[C:2](=[OH0:3])-[CH2:4][C:5](=[O:6])[O:7][C:8][C:9](-[OH])=[O:10]"

- name: reverse tetronate formation (type 3; spirotetronate)
  smarts: "[C:9]=[C:1]1[O:2][C:3](=[O:4])[C:5](-[O:6])=[C:7](-[OH:8])1>>([C:9]=[C:1]1[O:2][C:3](=[O:4])[C:5](-[O:6]).[OH][C:7](=[O:8])1)"

- name: reverse acetylation
  smarts: "[CH2:1]=[C:2](-[OH:3])[C:4](=[O:5])-[OH:6]>>[OH][CH2:1]-[C:2](-[OH:3])[C:4](=[O:5])-[OH:6]"

- name: reverse kirromycin-like substructure formation
  smarts: "[c:1]1([C:2](=[O:8])[*:9])[c:3](~[O:10])[c:4][c:5][n:6][c:7](~[O:11])1>>[CH0:3](=[O:10])(-[OH])[CH2:4][CH2:5][NH1:6][CH0:7](=[O:11])[CH2:1]([CH0:2](=[O:8])[*:9])"

- name: reverse beta-lactam formation
  smarts: "[N:1][C:2]1[C:3]2[S:4][C:5][C:6]([C:7])[N:8]2[C:9]1=[O:10]>>([N:1][C:2]1[C:3][S:4].[C:5][C:6]([C:7])[N:8][C:9]1=[O:10])"

- name: reverse DAOC-synthetase
  smarts: "[C:1][N:2][C:3]1[C:4]2[S:5][C:6][CH0:7]=[CH0:8]([C:9])[N:10]2[C:11]1=[O:12]>>[C:1][N:2][C:3]1[C:4]2[S:5][CH0:7]([CH3:6])[CH1:8]([C:9])[N:10]2[C:11]1=[O:12]"

- name: reverse salinosporamide-like substructure formation
  smarts: "[*:1][C:2]1[C:3](=[O:4])[N:5][C:6]([C:7](=[O:8])2)([*:9])[C:10]1([C:11])[O:12]2>>[*:1][C:2][C:3](=[O:4])[OH].[N:5][C:6]([C:7](=[O:8])[OH])([*:9]).[C:10]([C:11])(=[O])[OH:12]"

- name: reverse cyclization on lysine-like substructure
  smarts: "[C:1]1(-[N:2])[C:3][C:4][C:5][C:6][N:7]([OH:8])[C:9](=[O:10])1>>([C:1](-[N:2])([C:9](=[O:10])([OH]))[C:3][C:4][C:5][C:6][N:7]([OH0:8]))"

- name: open terminal aromatic ring
  smarts: "[c:1]1[c:2]([C:7](=[O:8])([OH:9]))[c:3]([C:10])[c:4][c:5][c:6]1>>([C:1]1=[C:2]([C:7](=[O:8])([OH:9])).[C:3]([C:10])=[C:4][C:5]=[C:6]1)"

- name: reverse spirocycle formation (type 1)
  smarts: "[#6:1]1(-[*:14])-[#8:6]-[#6:5]2(-[#8:7]-[#6:11](-[#6:13]-[*:12])-[#6:10]-[#6:9]-[#6:8]-2)-[#6:4]-[#6:3]-[#6:2]-1>>[#6:1](-[*:14])(-[#8:6])-[#6:2]-[#6:3]-[#6:4]-[#6:5](=[#8:7])-[#6:8]-[#6:9]-[#6:10]-[#6:11]=[#6:13]-[*:12]"

- name: reverse spirocycle formation (type 2)
  smarts: "[#6:1]1(-[*:13])-[#8:6]-[#6:5]2(-[#8:7]-[#6:10](-[#6:9]-[#6:8]-2)-[#6:12]-[*:11])-[#6:4]-[#6:3]-[#6:2]-1>>[#6:1](-[*:13])(-[#8:6])-[#6:2]-[#6:3]-[#6:4]-[#6:5](=[#8:7])-[#6:8]-[#6:9]-[#6:10]=[#6:12]-[*:11]"

- name: reversed cyclized hydroxy ornithine
  smarts: "[C:2]1[C:3](=[O:4])[NH0:5]([OH:7])[C:6][C:8][C:9]1>>[NH2:5]([OH:7])[C:6][C:8][C:9][C:2][C:3](=[O:4])[O]"

- name: reverse atrop-abyssomycin C like formation
  smarts: "[C;R:1]=[C;R:2][O;R:3][C;R:4][C;R:5][O:6]>>([C;R:1]=[C;R:2][OH1:3].[C;R:4]1[C;R:5][OH0:6]1)"

- name: reverse dies-alder (intramolecular)
  smarts: "[*:9][C:1]1-[C:2]=[C:3]-[C:4](-[C:5])-[C:6]-[C:7]1-[*:8]>>([*:9][C:1]=[C:2]-[C:3]=[C:4]-[C:5].[C:6]=[C:7]-[*:8])"

- name: reverse dies-alder (intermolecular)
  smarts: "[*:9][C:1]1-[C:2]=[C:3]-[C:4](-[C:5])-[C:6]-[C:7]1-[*:8]>>[*:9][C:1]=[C:2]-[C:3]=[C:4]-[C:5].[C:6]=[C:7]-[*:8]"

- name: reverse aryl amicoumacin-like
  smarts: "[C:1][c:2]1[c:3]([OH:9])[c:4][c:5][c:6][c:7]1[C:8]>>[C:1][C:2]=[C:3]([OH:9])[C:4]=[C:5][C:6]=[C:7][C:8]"

- name: reverse spiroborate
  smarts: "[O:1]1[C:2][C:3][O:4][B:5]12[O:6][C:7][C:8][O:9]2>>([OH1:1][C:2][C:3][OH1:4].[OH1:6][C:7][C:8][OH1:9]).[B:5]"

- name: reverse cremimycin-like substructure
  smarts: "[C:1]1[C:2][C:3]([O:4])[C:5](-,=[O:6])-,=[C:7]1>>[C:1]=[C:2][C:3][C:5](-[OH:6])-[C:7].[O:4]"
  props:
    references:
      - "DOI:10.1002/cbic.201300370"

- name: reversed NRP biosynthesis (alpha amino acid; intermolecular)
  smarts: "[C,c:2][C:3](=[O:4])[NH1:5][C,c;!$(C=O):6]>>[C,c:2][C:3](=[O:4])[O].[NH2:5][C,c:6]"
  allowed_in_bulk: true

- name: reversed NRP biosynthesis (alpha amino acid; intramolecular)
  smarts: "[C,c:2][C;R:3](=[O:4])[NH1;R:5][C,c;!$(C=O):6]>>([C,c:2][C:3](=[O:4])[O].[NH2:5][C,c:6])"
  allowed_in_bulk: true

- name: reverse NRP biosynthesis (aromatic nitrogen; intermolecular)
  smarts: "[*:1][C:2](=[O:3])[n:4]>>[*:1][C:2](=[O:3])[OH].[nH:4]"
  allowed_in_bulk: true

- name: reverse NRP biosynthesis (aromatic nitrogen; intramolecular)
  smarts: "[*:1][C:2](=[O:3])[n:4]>>([*:1][C:2](=[O:3])[OH].[nH:4])"
  allowed_in_bulk: true

- name: reverse urea bond formation (intermolecular)
  smarts: "[C:1][NH1:2][C:3](=[O:4])[NH1:5][C:6]>>[C:1][NH2:2].[NH2:5][C:6].[OH][C:3](=[O:4])[OH]"
  allowed_in_bulk: true

- name: reverse urea bond formation (intramolecular)
  smarts: "[C:1][NH1:2][C:3](=[O:4])[NH1:5][C:6]>>([C:1][NH2:2].[NH2:5][C:6]).[OH][C:3](=[O:4])[OH]"
  allowed_in_bulk: true

- name: reversed NRP biosynthesis (alpha amino acid; proline-like)
  smarts: "[*:1][C:2](=[O:3])[NH0:4][C:5][C:6](=[O:7])[OH:8]>>[*:1][C:2](=[O:3])[OH].[NH1:4][C:5][C:6](=[O:7])[OH:8]"
  allowed_in_bulk: true

- name: reversed NRP biosynthesis (beta amino acid)
  smarts: "[*:1][C:2](=[O:3])[NH1:4][C:5][CH2:6][C:7](=[O:8])[OH:9]>>[*:1][C:2](=[O:3])[OH].[NH2:4][C:5][CH2:6][C:7](=[O:8])[OH:9]"
  allowed_in_bulk: true

- name: break amide bond (hydroxamic)
  smarts: "[C:1][C:2](=[O:3])[N:4](-[OH:5])[C:6]>>[C:1][C:2](=[O:3])-[OH].[N:4](-[OH:5])[C:6]"
  allowed_in_bulk: true

- name: reversed polyketide synthesis (saturated)
  smarts: "[C,c:1][C;!R:2]-[C;!R:3]-[C:4](=[O:5])[OH:6]>>[C,c:1]C(=O)[OH].[OH][S][C:2]-[C:3]-[C:4](=[O:5])[OH:6]"

- name: reversed polyketide synthesis (saturated anhydride)
  smarts: "[C,c:1][C;!R:2]-[C;!R:3]-[C:4](=[O:5])[CH2:6][CH3:7]>>[C,c:1]C(=O)[OH].[OH][S][C:2]-[C:3]-[C:4](=[O:5])[OH].[C](=[O])([OH])[C:6][C:7]"

- name: reversed polyketide synthesis (unsaturated)
  smarts: "[C,c:1][C;!R:2]=[C;!R:3]-[C:4](=[O:5])[OH:6]>>[C,c:1]C(=O)[OH].[OH][S][C:2]=[C:3]-[C:4](=[O:5])[OH:6]"

- name: reversed polyketide synthesis (unsaturated and shifted type 1)
  smarts: "[C,c:1]=[C;!R:2]-[C;!R:3]-[C:4](=[O:5])[OH:6]>>[C,c:1]C(=O)[OH].[OH][S][C:2]=[C:3]-[C:4](=[O:5])[OH:6]"

- name: reversed polyketide synthesis (unsaturated and shifted type 2)
  smarts: "[C:7][C:1]=[C;!R:2]-[C&D4;!R:3]-[C:4](=[O:5])[OH:6]>>[C:7]=[C:1]C(=O)[OH].[OH][S][C:2]-[C:3]-[C:4](=[O:5])[OH:6]"

- name: reversed polyketide synthesis (shifted and late stage oxidation)
  smarts: "[C,c:1]=[C;!R:2]-[C;!R:3](-[OH:7])-[C:4](=[O:5])[OH:6]>>[C,c:1]C(=O)[OH].[OH][S][C:2]=[C:3]-[C:4](=[O:5])[OH:6].[O:7]"

- name: reverse polyketide unit carbonylation
  smarts: "[O:1][S:2][C:3]([C:4][C:5]([O:6])=[O:7])=[C:8][C:9]([O:10])=[O:11]>>[O:1][S:2][C:3]([C:4][C:5]([O:6])=[O:7])=O.[C:8][C:9]([O:10])=[O:11]"
```
