## Supplementary material for "RetroMol: Parsing a shared encoding from natural products and their biosynthetic gene clusters": RetroMol Supplemental Information: S5_default_matching_rules.yml

```
- name: A1
  smiles: "O=C(O)CC(=O)SO"
  pseudonyms: ["PK_A", "PK"]
- name: A1
  smiles: "O=C(O)C=C(O)SO"
  pseudonyms: ["PK_A", "PK"]

- name: A2^R
  smiles: "C[C@@H](C(SO)=O)C(O)=O"
  stereochemistry: true
  pseudonyms: ["A2", "PK_A", "PK"]
- name: A2^S
  smiles: "C[C@H](C(SO)=O)C(O)=O"
  stereochemistry: true
  pseudonyms: ["A2", "PK_A", "PK"]
- name: A2
  smiles: "CC(C(=O)O)C(=O)SO"
  pseudonyms: ["PK_A", "PK"]
- name: A2
  smiles: "CC(C(=O)O)=C(O)SO"
  pseudonyms: ["PK_A", "PK"]

- name: A3
  smiles: "CC(C)(C(=O)O)C(=O)SO"
  pseudonyms: ["PK_A", "PK"]

- name: A4^R
  smiles: "CC[C@@H](C(SO)=O)C(O)=O"
  stereochemistry: true
  pseudonyms: ["A4", "PK_A", "PK"]
- name: A4^S
  smiles: "CC[C@H](C(SO)=O)C(O)=O"
  stereochemistry: true
  pseudonyms: ["A4", "PK_A", "PK"]
- name: A4
  smiles: "CCC(C(=O)O)C(=O)SO"
  pseudonyms: ["PK_A", "PK"]

- name: A5^R
  smiles: "O=C([C@H](C(SO)=O)O)O"
  stereochemistry: true
  pseudonyms: ["A5", "PK_A", "PK"]
- name: A5^S
  smiles: "O=C([C@@H](C(SO)=O)O)O"
  stereochemistry: true
  pseudonyms: ["A5", "PK_A", "PK"]
- name: A5
  smiles: "O=C(O)C(O)C(=O)SO"
  pseudonyms: ["PK_A", "PK"]
- name: A5
  smiles: "O=C(C(O)=C(SO)O)O"
  pseudonyms: ["PK_A", "PK"]

- name: A6^R
  smiles: "C[C@@](C(O)=O)(C(SO)=O)O"
  stereochemistry: true
  pseudonyms: ["A6", "PK_A", "PK"]
- name: A6^S
  smiles: "C[C@](C(O)=O)(C(SO)=O)O"
  stereochemistry: true
  pseudonyms: ["A6", "PK_A", "PK"]
- name: A6
  smiles: "CC(O)(C(=O)O)C(=O)SO"
  pseudonyms: ["PK_A", "PK"]

- name: A7^R
  smiles: "O=C([C@H](C(SO)=O)CO)O"
  stereochemistry: true
  pseudonyms: ["A7", "PK_A", "PK"]
- name: A7^S
  smiles: "O=C([C@@H](C(SO)=O)CO)O"
  stereochemistry: true
  pseudonyms: ["A7", "PK_A", "PK"]
- name: A7
  smiles: "O=C(O)C(CO)C(=O)SO"
  pseudonyms: ["PK_A", "PK"]
- name: A7
  smiles: "O=C(O)C(CO)=C(O)SO"
  pseudonyms: ["PK_A", "PK"]

- name: A8
  smiles: "C=C(CC(=O)O)SO"
  pseudonyms: ["PK_A", "PK"]
- name: A8
  smiles: "CC(=CC(=O)O)SO"
  pseudonyms: ["PK_A", "PK"]

- name: A9^R
  smiles: "C=C([C@@H](C(O)=O)C)SO"
  stereochemistry: true
  pseudonyms: ["A9", "PK_A", "PK"]
- name: A9^S
  smiles: "C=C([C@H](C(O)=O)C)SO"
  stereochemistry: true
  pseudonyms: ["A9", "PK_A", "PK"]
- name: A9
  smiles: "C=C(SO)C(C)C(=O)O"
  pseudonyms: ["PK_A", "PK"]

- name: A10^R
  smiles: "C=C([C@@H](C(O)=O)O)SO"
  stereochemistry: true
  pseudonyms: ["A10", "PK_A", "PK"]
- name: A10^S
  smiles: "C=C([C@H](C(O)=O)O)SO"
  stereochemistry: true
  pseudonyms: ["A10", "PK_A", "PK"]
- name: A10
  smiles: "C=C(SO)C(O)C(=O)O"
  pseudonyms: ["PK_A", "PK"]

- name: A11
  smiles: "C=C(SO)C(=O)C(=O)O"
  pseudonyms: ["PK_A", "PK"]

- name: B^R1
  smiles: "O=C(C[C@@H](SO)O)O"
  stereochemistry: true
  pseudonyms: ["B1", "PK_B", "PK"]
- name: B^S1
  smiles: "O=C(C[C@H](SO)O)O"
  stereochemistry: true
  pseudonyms: ["B1", "PK_B", "PK"]
- name: B1
  smiles: "O=C(O)CC(O)SO"
  pseudonyms: ["PK_B", "PK"]

- name: B^R2^R
  smiles: "C[C@@H]([C@@H](SO)O)C(O)=O"
  stereochemistry: true
  pseudonyms: ["B2", "PK_B", "PK"]
- name: B^R2^S
  smiles: "C[C@H]([C@@H](SO)O)C(O)=O"
  stereochemistry: true
  pseudonyms: ["B2", "PK_B", "PK"]
- name: B^R2
  smiles: "CC([C@@H](SO)O)C(O)=O"
  stereochemistry: true
  pseudonyms: ["B2", "PK_B", "PK"]
- name: B^S2^R
  smiles: "C[C@@H]([C@H](SO)O)C(O)=O"
  stereochemistry: true
  pseudonyms: ["B2", "PK_B", "PK"]
- name: B^S2^S
  smiles: "C[C@H]([C@H](SO)O)C(O)=O"
  stereochemistry: true
  pseudonyms: ["B2", "PK_B", "PK"]
- name: B^S2
  smiles: "CC([C@H](SO)O)C(O)=O"
  stereochemistry: true
  pseudonyms: ["B2", "PK_B", "PK"]
- name: B2
  smiles: "CC(C(=O)O)C(O)SO"
  pseudonyms: ["PK_B", "PK"]

- name: B^R3
  smiles: "CC(C(O)=O)([C@@H](SO)O)C"
  stereochemistry: true
  pseudonyms: ["B3", "PK_B", "PK"]
- name: B^S3
  smiles: "CC(C(O)=O)([C@H](SO)O)C"
  stereochemistry: true
  pseudonyms: ["B3", "PK_B", "PK"]
- name: B3
  smiles: "CC(C)(C(=O)O)C(O)SO"
  pseudonyms: ["PK_B", "PK"]

- name: B^R4^R
  smiles: "CC[C@@H]([C@@H](SO)O)C(O)=O"
  stereochemistry: true
  pseudonyms: ["B4", "PK_B", "PK"]
- name: B^R4^S
  smiles: "CC[C@H]([C@@H](SO)O)C(O)=O"
  stereochemistry: true
  pseudonyms: ["B4", "PK_B", "PK"]
- name: B^S4^R
  smiles: "CC[C@@H]([C@H](SO)O)C(O)=O"
  stereochemistry: true
  pseudonyms: ["B4", "PK_B", "PK"]
- name: B^S4^S
  smiles: "CC[C@H]([C@H](SO)O)C(O)=O"
  stereochemistry: true
  pseudonyms: ["B4", "PK_B", "PK"]
- name: B4
  smiles: "CCC(C(=O)O)C(O)SO"
  pseudonyms: ["PK_B", "PK"]

- name: B^R5^R
  smiles: "O=C([C@H]([C@@H](SO)O)O)O"
  stereochemistry: true
  pseudonyms: ["B5", "PK_B", "PK"]
- name: B^R5^S
  smiles: "O=C([C@@H]([C@@H](SO)O)O)O"
  stereochemistry: true
  pseudonyms: ["B5", "PK_B", "PK"]
- name: B^S5^R
  smiles: "O=C([C@H]([C@H](SO)O)O)O"
  stereochemistry: true
  pseudonyms: ["B5", "PK_B", "PK"]
- name: B^S5^S
  smiles: "O=C([C@@H]([C@H](SO)O)O)O"
  stereochemistry: true
  pseudonyms: ["B5", "PK_B", "PK"]
- name: B5
  smiles: "O=C(O)C(O)C(O)SO"
  pseudonyms: ["PK_B", "PK"]

- name: B^R6^R
  smiles: "C[C@@](C(O)=O)([C@@H](SO)O)O"
  stereochemistry: true
  pseudonyms: ["B6", "PK_B", "PK"]
- name: B^R6^S
  smiles: "C[C@](C(O)=O)([C@@H](SO)O)O"
  stereochemistry: true
  pseudonyms: ["B6", "PK_B", "PK"]
- name: B^S6^R
  smiles: "C[C@@](C(O)=O)([C@H](SO)O)O"
  stereochemistry: true
  pseudonyms: ["B6", "PK_B", "PK"]
- name: B^S6^S
  smiles: "C[C@](C(O)=O)([C@H](SO)O)O"
  stereochemistry: true
  pseudonyms: ["B6", "PK_B", "PK"]
- name: B6
  smiles: "CC(O)(C(=O)O)C(O)SO"
  pseudonyms: ["PK_B", "PK"]

- name: B^R7^R
  smiles: "O=C([C@H]([C@@H](SO)O)CO)O"
  stereochemistry: true
  pseudonyms: ["B7", "PK_B", "PK"]
- name: B^R7^S
  smiles: "O=C([C@@H]([C@@H](SO)O)CO)O"
  stereochemistry: true
  pseudonyms: ["B7", "PK_B", "PK"]
- name: B^S7^R
  smiles: "O=C([C@H]([C@H](SO)O)CO)O"
  stereochemistry: true
  pseudonyms: ["B7", "PK_B", "PK"]
- name: B^S7^S
  smiles: "O=C([C@@H]([C@H](SO)O)CO)O"
  stereochemistry: true
  pseudonyms: ["B7", "PK_B", "PK"]
- name: B7
  smiles: "O=C(O)C(CO)C(O)SO"
  pseudonyms: ["PK_B", "PK"]

- name: B^R8
  smiles: "C[C@](CC(O)=O)(SO)O"
  stereochemistry: true
  pseudonyms: ["B8", "PK_B", "PK"]
- name: B^S8
  smiles: "C[C@@](CC(O)=O)(SO)O"
  stereochemistry: true
  pseudonyms: ["B8", "PK_B", "PK"]
- name: B8
  smiles: "CC(O)(CC(=O)O)SO"
  pseudonyms: ["PK_B", "PK"]

- name: B^R9^R
  smiles: "C[C@@H]([C@](O)(SO)C)C(O)=O"
  stereochemistry: true
  pseudonyms: ["B9", "PK_B", "PK"]
- name: B^R9^S
  smiles: "C[C@H]([C@](O)(SO)C)C(O)=O"
  stereochemistry: true
  pseudonyms: ["B9", "PK_B", "PK"]
- name: B^S9^R
  smiles: "C[C@@H]([C@@](O)(SO)C)C(O)=O"
  stereochemistry: true
  pseudonyms: ["B9", "PK_B", "PK"]
- name: B^S9^S
  smiles: "C[C@H]([C@@](O)(SO)C)C(O)=O"
  stereochemistry: true
  pseudonyms: ["B9", "PK_B", "PK"]
- name: B9
  smiles: "CC(C(=O)O)C(C)(O)SO"
  pseudonyms: ["PK_B", "PK"]

- name: B^R10^R
  smiles: "C[C@@](SO)([C@@H](C(O)=O)O)O"
  stereochemistry: true
  pseudonyms: ["B10", "PK_B", "PK"]
- name: B^R10^S
  smiles: "C[C@@](SO)([C@H](C(O)=O)O)O"
  stereochemistry: true
  pseudonyms: ["B10", "PK_B", "PK"]
- name: B^S10^R
  smiles: "C[C@](SO)([C@@H](C(O)=O)O)O"
  stereochemistry: true
  pseudonyms: ["B10", "PK_B", "PK"]
- name: B^S10^S
  smiles: "C[C@](SO)([C@H](C(O)=O)O)O"
  stereochemistry: true
  pseudonyms: ["B10", "PK_B", "PK"]
- name: B10
  smiles: "CC(O)(SO)C(O)C(=O)O"
  pseudonyms: ["PK_B", "PK"]

- name: B11
  smiles: "O=C(O)C(=O)C(O)SO"
  pseudonyms: ["PK_B", "PK"]

- name: B^R12^R
  smiles: "N[C@@H]([C@@H](SO)O)C(O)=O"
  stereochemistry: true
  pseudonyms: ["B12", "PK_B", "PK"]
- name: B^R12^S
  smiles: "N[C@H]([C@@H](SO)O)C(O)=O"
  stereochemistry: true
  pseudonyms: ["B12", "PK_B", "PK"]
- name: B^S12^R
  smiles: "N[C@@H]([C@H](SO)O)C(O)=O"
  stereochemistry: true
  pseudonyms: ["B12", "PK_B", "PK"]
- name: B^S12^S
  smiles: "N[C@H]([C@H](SO)O)C(O)=O"
  stereochemistry: true
  pseudonyms: ["B12", "PK_B", "PK"]
- name: B12
  smiles: "NC(C(=O)O)C(O)SO"
  pseudonyms: ["PK_B", "PK"]

- name: B^R13
  smiles: "OS[C@H](C(C(O)=O)C(O)=O)O"
  stereochemistry: true
  pseudonyms: ["B13", "PK_B", "PK"]
- name: B^S13
  smiles: "OS[C@@H](C(C(O)=O)C(O)=O)O"
  stereochemistry: true
  pseudonyms: ["B13", "PK_B", "PK"]
- name: B13
  smiles: "OSC(O)C(C(=O)O)C(=O)O"
  pseudonyms: ["PK_B", "PK"]

- name: C1
  smiles: "O=C(O)C=CSO"
  pseudonyms: ["PK_C", "PK"]

- name: C2
  smiles: "CC(=CSO)C(=O)O"
  pseudonyms: ["PK_C", "PK"]

- name: C4
  smiles: "CCC(=CSO)C(=O)O"
  pseudonyms: ["PK_C", "PK"]

- name: C7
  smiles: "O=C(O)C(=CSO)CO"
  pseudonyms: ["PK_C", "PK"]

- name: C13
  smiles: "O=C(C(C(O)=O)=CSO)O"
  pseudonyms: ["PK_C", "PK"]

- name: D1
  smiles: "O=C(O)CCSO"
  pseudonyms: ["PK_D", "PK"]

- name: D2^R
  smiles: "C[C@H](C(O)=O)CSO"
  stereochemistry: true
  pseudonyms: ["D2", "PK_D", "PK"]
- name: D2^S
  smiles: "C[C@@H](C(O)=O)CSO"
  stereochemistry: true
  pseudonyms: ["D2", "PK_D", "PK"]
- name: D2
  smiles: "CC(CSO)C(=O)O"
  pseudonyms: ["PK_D", "PK"]

- name: D3
  smiles: "CC(C)(CSO)C(=O)O"
  pseudonyms: ["PK_D", "PK"]

- name: D4^R
  smiles: "CC[C@H](C(O)=O)CSO"
  stereochemistry: true
  pseudonyms: ["D4", "PK_D", "PK"]
- name: D4^S
  smiles: "CC[C@@H](C(O)=O)CSO"
  stereochemistry: true
  pseudonyms: ["D4", "PK_D", "PK"]
- name: D4
  smiles: "CCC(CSO)C(=O)O"
  pseudonyms: ["PK_D", "PK"]

- name: D5^R
  smiles: "O=C([C@H](CSO)O)O"
  pseudonyms: ["D5", "PK_D", "PK"]
  stereochemistry: true
- name: D5^S
  smiles: "O=C([C@@H](CSO)O)O"
  stereochemistry: true
  pseudonyms: ["D5", "PK_D", "PK"]
- name: D5
  smiles: "O=C(O)C(O)CSO"
  pseudonyms: ["PK_D", "PK"]

- name: D6^R
  smiles: "C[C@](CSO)(C(O)=O)O"
  stereochemistry: true
  pseudonyms: ["D6", "PK_D", "PK"]
- name: D6^S
  smiles: "C[C@@](CSO)(C(O)=O)O"
  stereochemistry: true
  pseudonyms: ["D6", "PK_D", "PK"]
- name: D6
  smiles: "CC(O)(CSO)C(=O)O"
  pseudonyms: ["PK_D", "PK"]

- name: D7^R
  smiles: "O=C([C@H](CSO)CO)O"
  stereochemistry: true
  pseudonyms: ["D7", "PK_D", "PK"]
- name: D7^S
  smiles: "O=C([C@@H](CSO)CO)O"
  stereochemistry: true
  pseudonyms: ["D7", "PK_D", "PK"]
- name: D7
  smiles: "O=C(O)C(CO)CSO"
  pseudonyms: ["PK_D", "PK"]

- name: D^R8
  smiles: "C[C@@H](SO)CC(O)=O"
  stereochemistry: true
  pseudonyms: ["D8", "PK_D", "PK"]
- name: D^S8
  smiles: "C[C@H](SO)CC(O)=O"
  stereochemistry: true
  pseudonyms: ["D8", "PK_D", "PK"]
- name: D8
  smiles: "CC(CC(=O)O)SO"
  pseudonyms: ["PK_D", "PK"]

- name: D^R10^R
  smiles: "C[C@H]([C@@H](C(O)=O)O)SO"
  stereochemistry: true
  pseudonyms: ["D10", "PK_D", "PK"]
- name: D^R10^S
  smiles: "C[C@H]([C@H](C(O)=O)O)SO"
  stereochemistry: true
  pseudonyms: ["D10", "PK_D", "PK"]
- name: D^S10^R
  smiles: "C[C@@H]([C@@H](C(O)=O)O)SO"
  stereochemistry: true
  pseudonyms: ["D10", "PK_D", "PK"]
- name: D^S10^S
  smiles: "C[C@@H]([C@H](C(O)=O)O)SO"
  stereochemistry: true
  pseudonyms: ["D10", "PK_D", "PK"]
- name: D10
  smiles: "CC(SO)C(O)C(=O)O"
  pseudonyms: ["PK_D", "PK"]

- name: D11
  smiles: "O=C(O)C(=O)CSO"
  pseudonyms: ["PK_D", "PK"]
- name: D11
  smiles: "O=C(O)C(O)=CSO"
  pseudonyms: ["PK_D", "PK"]

- name: D14
  smiles: "CC(O)C(CSO)C(=O)O"
  pseudonyms: ["PK_D", "PK"]

- name: D15
  smiles: "C=C(C(O)=O)CSO"
  pseudonyms: ["PK_D", "PK"]

- name: D16^R
  smiles: "OSC[C@@H](C(O)=O)CC=O"
  stereochemistry: true
  pseudonyms: ["D16", "PK_D", "PK"]
- name: D16^S
  smiles: "OSC[C@H](C(O)=O)CC=O"
  stereochemistry: true
  pseudonyms: ["D16", "PK_D", "PK"]
- name: D16
  smiles: "OSCC(CC=O)C(=O)O"
  pseudonyms: ["PK_D", "PK"]

- name: D^R17
  smiles: "O=C(O)C([C@H](SO)C)=O"
  stereochemistry: true
  pseudonyms: ["D17", "PK_D", "PK"]
- name: D^S17
  smiles: "O=C(O)C([C@@H](SO)C)=O"
  stereochemistry: true
  pseudonyms: ["D17", "PK_D", "PK"]
- name: D17
  smiles: "O=C(C(C(C)SO)=O)O"
  pseudonyms: ["PK_D", "PK"]
- name: D17
  smiles: "O=C(C(O)=C(C)SO)O"
  pseudonyms: ["PK_D", "PK"]

- name: A
  smiles: "OSC(N)CC(=O)O"
  pseudonyms: ["PK"]

- name: A
  smiles: "OSC(N)C(C)C(=O)O"
  pseudonyms: ["PK"]

- name: A
  smiles: "OSC(N)C(C)(C)C(=O)O"
  pseudonyms: ["PK"]

- name: A
  smiles: "OSC(N)C(CC)C(=O)O"
  pseudonyms: ["PK"]

- name: A
  smiles: "OSC(N)C(C)(O)C(=O)O"
  pseudonyms: ["PK"]

- name: A
  smiles: "NC(C(O)=O)CSO"
  pseudonyms: ["PK"]

- name: A
  smiles: "NC(C(C(O)=O)O)SO"
  pseudonyms: ["PK"]

- name: A
  smiles: "O=C(C(C(C(O)=O)SO)O)O"
  pseudonyms: ["PK"]

- name: methylation
  smiles: "C"

- name: oxidation
  smiles: "O"

- name: amination
  smiles: "N"

- name: cyanide
  smiles: "C#N"

- name: boronation
  smiles: "B"

- name: phosphonylation
  smiles: "OP(O)(O)O"

- name: fluorination
  smiles: "F"
  pseudonyms: ["halogenation"]

- name: chlorination
  smiles: "Cl"
  pseudonyms: ["halogenation"]

- name: bromination
  smiles: "Br"
  pseudonyms: ["halogenation"]

- name: iodination
  smiles: "I"
  pseudonyms: ["halogenation"]

- name: methanol
  smiles: "CO"

- name: ethanol
  smiles: "CCO"

- name: propanol
  smiles: "CCCO"

- name: formaldehyde
  smiles: "C=O"

- name: carbamic acid
  smiles: "NC(=O)O"

- name: carbonic acid
  smiles: "OC(=O)O"

- name: acetic acid
  smiles: "CC(=O)O"

- name: glyceric acid
  smiles: "OC(CO)C(O)=O"

- name: propanoic acid
  smiles: "CCC(O)=O"

- name: glycosylation
  smiles: "CC1CC(N)C(O)C(O)O1"

- name: glycosylation
  smiles: "CC1OC(O)CC(N)C1O"

- name: glycosylation
  smiles: "CC1OC(O)CC(C)(O)C1O"

- name: glycosylation
  smiles: "CC1CC(O)C(O)C(O)O1"

- name: glycosylation
  smiles: "NC1C(O)OC(O)C(O)C1O"

- name: glycosylation
  smiles: "OC1COC(O)C(O)C1O"

- name: glycosylation
  smiles: "OCC1OC(O)C(O)C(O)C1O"

- name: glycosylation
  smiles: "CC1OC(O)CCC1O"

- name: glycosylation
  smiles: "CC1OC(O)CC(O)C1O"

- name: glycosylation
  smiles: "CC1OC(O)C(O)C(O)C1O"

- name: glycosylation
  smiles: "CC1OC(O)C(O)C(N)C1O"

- name: glycosylation
  smiles: "CC(C1(C(C(C(OC1C)O)O)O)O)O"

- name: glycosylation
  smiles: "CC(C1(O)C(O)C(O)C(O)OC1C)=O"

- name: glycosylation
  smiles: "NC1C(C(C(OC1O)CO)O)O"

- name: glycosylation
  smiles: "NC1COC(CC1O)O"

- name: glycosylation
  smiles: "OC1C(O)C(N)C(C)OC1O"

- name: glycosylation
  smiles: "CC1OC(CC(C1S)O)O"

- name: glycosylation
  smiles: "CC1(N)CC(O)OC(C)C1O"

- name: glycosylation
  smiles: "NC1C(O)C(O)C(O)C(N)C1"

- name: glycosylation
  smiles: "O=CC1(O)C(CO)OC(O)C1O"

# Tautomers
- name: glycosylation
  smiles: "NC(N)=NC1C(O)C(O)C(O)C(N=C(N)N)C1O"
- name: glycosylation
  smiles: "N=C(N)NC1C(O)C(NC(=N)N)C(O)C(O)C1O"

- name: glycosylation
  smiles: "OCC1OC(O)C(O)C1O"

- name: glycosylation
  smiles: "NCC1OC(C(C(C1O)O)N)O"

- name: glycosylation
  smiles: "OC1CC(C)(OC)C(O)C(C)O1"

- name: glycosylation
  smiles: "OC1C(O)C(N(C)C)CC(C)O1"

# Tautomers
- name: arginine
  smiles: "NC(N)=NCCCC(N)C(=O)O"
- name: arginine
  smiles: "N=C(N)NCCCC(N)C(=O)O"

- name: histidine
  smiles: "NC(Cc1cnc[nH]1)C(=O)O"

- name: 3-hydroxy histdine
  smiles: "NC(C(c1[nH]cnc1)O)C(O)=O"

- name: lysine
  smiles: "NCCCCC(N)C(=O)O"

- name: aspartic acid
  smiles: "NC(CC(=O)O)C(=O)O"

- name: glutamic acid
  smiles: "NC(CCC(=O)O)C(=O)O"

- name: serine
  smiles: "NC(CO)C(=O)O"

- name: threonine
  smiles: "CC(O)C(N)C(=O)O"

- name: asparagine
  smiles: "NC(=O)CC(N)C(=O)O"

- name: glutamine
  smiles: "NC(=O)CCC(N)C(=O)O"

- name: glycine
  smiles: "NCC(=O)O"

- name: proline
  smiles: "O=C(O)C1CCCN1"

- name: cysteine
  smiles: "NC(CS)C(=O)O"

- name: leucine
  smiles: "CC(C)CC(N)C(=O)O"

- name: isoleucine
  smiles: "CCC(C)C(N)C(=O)O"

- name: alanine
  smiles: "CC(N)C(=O)O"

- name: valine
  smiles: "CC(C)C(N)C(=O)O"

- name: methionine
  smiles: "CSCCC(C(=O)O)N"

- name: phenylalanine
  smiles: "NC(Cc1ccccc1)C(=O)O"

- name: tryptophan
  smiles: "NC(Cc1c[nH]c2ccccc12)C(=O)O"

- name: tyrosine
  smiles: "NC(Cc1ccc(O)cc1)C(=O)O"

- name: 3,4-dihydroxyphenylalanine
  smiles: "NC(C(O)=O)Cc1c(O)cc(O)cc1"

- name: kynurenine
  smiles: "Nc1ccccc1C(=O)C[C@H](N)C(=O)O"

- name: ornithine
  smiles: "[NH2]CCCC([NH2])C(=O)O"

- name: ornithine alkaloid
  smiles: "NC(NCCCCO)=N"

- name: bicyclic guanidine alkaloid
  smiles: "CCCCCCCCCC1N=C(N2CCCC2=C1C(O)=O)N"

- name: ornithine alkaloid
  smiles: "NC(N)=NCCCCO"

- name: dehydrobutyrine
  smiles: "CC=C(C(O)=O)N"

- name: dehydrohomoserine
  smiles: "NC(C(O)=O)C=CO"

- name: N6-formyl-N6-hydroxylysine
  smiles: "NC(CCCCN(O)C=O)C(O)=O"

- name: 2,3-diaminopropionate
  smiles: "NCC(N)C(=O)O"

- name: 3-hydroxyquinaldic acid
  smiles: "O=C(O)c1nc2ccccc2cc1O"

- name: beta-alanine
  smiles: "NCCC(=O)O"

- name: salicylic acid
  smiles: "O=C(O)c1ccccc1O"

- name: 2,3-dihydroxybenzoic acid
  smiles: "O=C(O)c1cccc(O)c1O"

- name: 3,5-dihydroxybenzoic acid
  smiles: "O=C(c1cc(O)cc(O)c1)O"

- name: isobutyric acid
  smiles: "CC(C)C(=O)O"

- name: lactic acid
  smiles: "CC(O)C(=O)O"

- name: 2-methyl lactic acid 
  smiles: "CC(C)(O)C(=O)O"

- name: 2,2-dimethylpropanoic acid
  smiles: "CC(C)(C)C(=O)O"
  terminal: false

- name: butanoic acid
  smiles: "CCCC(O)=O"
  terminal: false

- name: 3-methylbutanoic acid
  smiles: "CC(CC(O)=O)C"
  terminal: false

- name: 4-amino-2-hydroxybutanoic acid
  smiles: "NCCC(C(O)=O)O"
  terminal: false

- name: 3-butynoic acid
  smiles: "C#CCC(=O)O"
  terminal: false

- name: pentanoic acid
  smiles: "CCCCC(O)=O"
  terminal: false

- name: hexanoic acid
  smiles: "CCCCCC(O)=O"
  terminal: false

- name: heptanoic acid
  smiles: "CCCCCCC(O)=O"
  terminal: false

- name: octanoic acid
  smiles: "CCCCCCCC(O)=O"
  terminal: false

- name: nonanoic acid
  smiles: "CCCCCCCCC(O)=O"
  terminal: false

- name: 2-hydroxy-3-aminononanoic acid
  smiles: "CCCCCCCC(N)C(O)C(=O)O"
  terminal: false

- name: decanoic acid
  smiles: "CCCCCCCCCC(O)=O"
  terminal: false

- name: dec-3-enoic acid
  smiles: "CCCCCCC=CCC(=O)O"
  terminal: false

- name: 3-hydroxydecanoic acid
  smiles: "CCCCCCCC(O)CC(=O)O"
  terminal: false

- name: dodecanoic acid
  smiles: "CCCCCCCCCCCC(=O)O"
  terminal: false

- name: tridecanoic acid
  smiles: "CCCCCCCCCCCCC(=O)O"
  terminal: false

- name: 3-hydroxytridecanoic acid
  smiles: "O=C(O)CC(O)CCCCCCCCCC"
  terminal: false

- name: tetradecanoic acid
  smiles: "CCCCCCCCCCCCCC(=O)O"
  terminal: false

- name: tetradec-2-enoic acid
  smiles: "CCCCCCCCCCCC=CC(=O)O"
  terminal: false

- name: pentadecanoic acid
  smiles: "CCCCCCCCCCCCCCC(=O)O"
  terminal: false

- name: hexadecanoic acid
  smiles: "CCCCCCCCCCCCCCCC(=O)O"
  terminal: false

- name: 2,3-dihydroxyhexadecanoic acid
  smiles: "CCCCCCCCCCCCCC(C(C(=O)O)O)O"
  terminal: false

- name: 3-hydroxy-2,2-dimethyldecanoic acid
  smiles: "CCCCCCCC(C(C)(C(O)=O)C)O"
  terminal: false

- name: trans-2-hexanoic acid
  smiles: "CCCC=CC(O)=O"
  terminal: false

- name: orsellinic acid
  smiles: "O=C(O)c1c(O)cc(O)cc1C"
  terminal: false

- name: 6-methylsalicylic acid
  smiles: "CC1=C(C(=CC=C1)O)C(=O)O"
  terminal: false

- name: 2-methylprop-2-enoic acid
  smiles: "C=C(C)C(=O)O"

- name: cyclopentane-1,2-dicarboxylic acid
  smiles: "O=C(O)C1CCCC1C(=O)O"

- name: furan-3-carboxylic acid
  smiles: "O=C(O)c1ccoc1"

- name: benzoic acid
  smiles: "O=C(O)c1ccccc1"

- name: 2-phenylacetic acid
  smiles: "O=C(O)Cc1ccccc1"

- name: 3-amino-5-hydroxybenzoic acid
  smiles: "Nc1cc(C(O)=O)cc(O)c1"

- name: 2-(2,6-dioxopiperidin-4-yl)acetic acid
  smiles: "O=C1NC(CC(CC(O)=O)C1)=O"

- name: artificial amino acid harzianic acid
  smiles: "NC(C(O)=O)CC(C(O)=O)(C(C)C)O"

- name: 4-hydroxyphenylglycine
  smiles: "NC(C(=O)O)c1ccc(O)cc1"

- name: 2,4-diaminobytyric acid
  smiles: "NCCC(N)C(=O)O"

- name: beta-hydroxytyrosine
  smiles: "NC(C(=O)O)C(O)c1ccc(O)cc1"

- name: 3,5-dihydroxyphenylglycine
  smiles: "NC(C(=O)O)c1cc(O)cc(O)c1"

- name: pipecolic acid
  smiles: "O=C(O)C1CCCCN1"

- name: N5-formyl-N5-hydroxyornithine
  smiles: "NC(CCCN(O)C=O)C(=O)O"

- name: N5-hydroxyornithine
  smiles: "NC(CCCNO)C(=O)O"

- name: anthranilic acid
  smiles: "Nc1ccccc1C(=O)O"

- name: 2-amino-3-hydroxy-4-(4-nitrophenyl)butanoic acid
  smiles: "C1=CC(=CC=C1CC(C(C(=O)O)N)O)[N+](=O)[O-]"

- name: 2,6-diamino-57-dihydroxy-heptanoic acid
  smiles: "C(CC(C(=O)O)N)C(C(CO)N)O"

- name: 4-methylhex-2-enoic acid
  smiles: "CCC(C)C=CC(=O)O"

- name: 2-amino-4-decenoic acid
  smiles: "CCCCCC=CCC(C(=O)O)N"

- name: 1-(1,1-dimethylallyl)-tryptophan
  smiles: "CC(C)(C=C)N1C=C(C2=CC=CC=C21)CC(C(=O)O)N"

- name: 1-aminocyclopropane-1-carboxylic acid
  smiles: "C(O)(=O)C1(CC1)(N)"

- name: 1-pyrroline-5-carboxylic acid
  smiles: "O=C(O)C1N=CCC1"

- name: 10,14-dimethyloctadecanoic acid
  smiles: "OC(CCCCCCCCC(C)CCCC(C)CCCC)=O"

- name: 2,3-diaminobutyric acid
  smiles: "NC(C)C(C(=O)O)N"

- name: 2,3-dihydroxy-para-aminobenzoic acid
  smiles: "C1=CC(=C(C(=C(N)1)O)O)C(=O)O"

- name: 2,4-dihydroxypentanoic acid
  smiles: "CC(CC(C(=O)O)O)O"

- name: 2-(1-methylcyclopropyl)-glycine
  smiles: "CC1(CC1)C(C(=O)O)N"

- name: 2-amino-3,5-dimethyl-4-hexenoic Acid
  smiles: "CC(C=C(C)C)C(C(=O)O)N"

- name: 2-amino-3-hydroxycyclopent-2-enone
  smiles: "C1CC(=O)C(=C1O)N"

- name: 2-amino-6-hydroxy-4-methyl-8-oxodecanoic acid
  smiles: "CCC(=O)CC(CC(C)CC(C(=O)O)N)O"

- name: 2-aminoadipic acid
  smiles: "C(CC(C(=O)O)N)CC(=O)O"

- name: 2-aminobutyric acid
  smiles: "CCC(C(=O)O)N"

- name: 2-aminoisobutyric acid
  smiles: "O=C(O)C(N)(C)C"

- name: 2-carboxy-6-hydroxyoctahydroindole
  smiles: "N1[C@H](C(=O)O)CC2CCC(O)CC12"

- name: 2-chloro-3,5-dihydroxy-4-methylphenylglycine
  smiles: "CC1=C(O)C(Cl)=C(C=C(O)1)C(C(=O)O)N"

- name: 2-chlorobenzoic acid
  smiles: "C1=CC=C(C(=C1)C(=O)O)Cl"

- name: 2-hydroxy-4-methylpentanoic acid
  smiles: "CC(C)CC(C(=O)O)O"

- name: 2-hydroxypent-4-enoic acid
  smiles: "C=CCC(C(=O)O)O"

- name: 2-ketoglutaric acid
  smiles: "C(CC(=O)O)C(=O)C(=O)O"

- name: 2-ketoisocaproic acid
  smiles: "O=C(C(=O)O)CC(C)C"

- name: 2-ketoisovaleric acid
  smiles: "O=C(C(=O)O)C(C)C"

- name: 2-methylserine
  smiles: "CC(CO)(C(=O)O)N"

- name: 2-sulfamoylacetic acid
  smiles: "C(C(=O)O)S(=O)(=O)N"

- name: 2-hydroxy-3-methylpentanoic acid
  smiles: "CCC(C)C(C(=O)O)O"

- name: 2-amino-4-hexenoic acid
  smiles: "CC=CCC(C(=O)O)N"

- name: 2-amino-8-oxodecanoic acid
  smiles: "CCC(=O)CCCCCC(C(=O)O)N"

- name: 2-amino-9,10-epoxy-8-oxodecanoic acid
  smiles: "C1C(O1)C(=O)CCCCCC(C(=O)O)N"

- name: 2-aminodecanoic acid
  smiles: "CCCCCCCCC(C(=O)O)N"

- name: 2-aminododecanoic acid
  smiles: "CCCCCCCCCCC(C(=O)O)N"

- name: 2-aminooctanoic acid
  smiles: "CCCCCCC(C(=O)O)N"

- name: 2-hydroxyisovaleric acid
  smiles: "CC(C)C(C(=O)O)O"

- name: 2-methyl-3-oxobutyrine
  smiles: "CC(=O)C(C)(N)C(=O)O"

- name: 3,3-dihomo-4-methoxytyrosine
  smiles: "NC(CCCC1=CC=C(OC)C=C1)C(=O)O"

- name: 3,3-dihomophenylalanine
  smiles: "NC(CCCC1=CC=CC=C1)C(=O)O"

- name: 3,3-dihomotyrosine
  smiles: "NC(CCCC1=CC=C(O)C=C1)C(=O)O"

- name: 3,4-dehydrolysine
  smiles: "C(CCN)=CC(C(=O)O)N"

- name: 3,4-dihydroxybenzoic acid
  smiles: "C1=CC(=C(C=C1C(=O)O)O)O"

- name: 3,5-dichloro-4-hydroxyphenylglycine
  smiles: "C1=C(Cl)C(=C(Cl)C=C1C(C(=O)O)N)O"

- name: 3-(2-nitrocyclopropylalanine)
  smiles: "C1C(C1[N+](=O)[O-])CC(C(=O)O)N"

- name: 3-(3ridyl)-alanine
  smiles: "C1=CC(=CN=C1)CC(C(=O)O)N"

- name: 3-amino-2,4-dihydroxybenzoic acid
  smiles: "C1=CC(=C(C(=C1C(=O)O)O)N)O"

- name: 3-amino-4-hydroxybenzoic acid
  smiles: "C1=CC(=C(C=C1C(=O)O)N)O"

- name: 3-amino-6-hydroxy-2-pridone
  smiles: "C1CC(NC(=O)C1N)O"

- name: 3-chlorotyrosine
  smiles: "C1=C(Cl)C(=CC=C1CC(C(=O)O)N)O"

- name: 3-hydroxy-4-methylproline
  smiles: "CC1C(O)C(NC1)C(=O)O"

- name: 3-hydroxy-O-methyl-5-methyltyrosine
  smiles: "C1=C(O)C(=C(C)C=C1CC(C(=O)O)N)OC"

- name: 3-hydroxy-O-methyltyrosine
  smiles: "C1=C(O)C(=CC=C1CC(C(=O)O)N)OC"

- name: 3-hydroxy-para-aminobenzoic acid
  smiles: "C1=CC(=C(C=C1C(=O)O)O)N"

- name: 3-hydroxyaspartic acid
  smiles: "NC(C(C(=O)O)O)(C(=O)O)"

- name: 2-hydroxyglycine
  smiles: "NC(C(O)=O)O"

- name: 3-hydroxyglutamine
  smiles: "C(C(C(C(=O)O)N)O)C(=O)N"

- name: 3-hydroxyglutamic acid
  smiles: "NC(C(CC(O)=O)O)C(O)=O"

- name: 3-hydroxykynurenine
  smiles: "C1=CC(=C(C(=C1)O)N)C(=O)CC(C(=O)O)N"

- name: 3-hydroxyleucine
  smiles: "CC(C)C(C(C(=O)O)N)O"

- name: 3-hydroxypicolinic acid
  smiles: "C1=CC(=C(N=C1)C(=O)O)O"

- name: 3-hydroxytyrosine
  smiles: "C1=CC(=C(C=C1CC(C(=O)O)N)O)O"

- name: 3-hydroxyvaline
  smiles: "CC(O)(C)C(C(=O)O)N"

- name: 3-methoxyanthranilic acid
  smiles: "COC1=CC=CC(=C1N)C(=O)O"

- name: 3-methoxyaspartic acid
  smiles: "NC(C(C(=O)O)OC)(C(=O)O)"

- name: 3-methylasparagine
  smiles: "CC(C(C(=O)O)N)C(=O)N"

- name: 3-methylaspartic acid
  smiles: "CC(C(C(=O)O)N)C(=O)O"

- name: 3-methylglutamic acid
  smiles: "CC(CC(=O)O)C(C(=O)O)N"

- name: 3-methylleucine
  smiles: "CC(C)C(C)C(C(=O)O)N"

- name: 3-nitrotyrosine
  smiles: "C1=CC(=C(C=C1CC(C(=O)O)N)[N+](=O)[O-])O"

- name: 3-chloroproline
  smiles: "C1C(Cl)C(NC1)C(=O)O"

- name: 3-hydroxy-2,4-diaminobutyric acid
  smiles: "NCC(O)C(C(=O)O)N"

- name: 3-hydroxyasparagine
  smiles: "NC(C(O)=O)C(O)C(N)=O"

- name: 3-hydroxyhomotyrosine
  smiles: "C1=CC(=CC=C1CC(C(C(=O)O)N)O)O"

- name: 3-methylbeta-alanine
  smiles: "NCC(C)C(=O)O"

- name: 3,4-dichloroproline
  smiles: "ClC1C(Cl)C(NC1)C(=O)O"

- name: 3,4-dihydroxyhomotyrosine
  smiles: "C1=CC(=CC=C1C(O)C(C(C(=O)O)N)O)O"

- name: 3-aminobutyric acid
  smiles: "CC(CC(=O)O)N"

- name: 3-cyclohex-2-enylalanine
  smiles: "C1C=CC(CC1)CC(C(=O)O)N"

- name: 3-hydroxy-4-methyloctanoic acid
  smiles: "CCCCC(C(CC(O)=O)O)C"

- name: 3-hydroxy-6-chlorohistidine
  smiles: "C1=C(NC(Cl)=N1)C(C(C(=O)O)N)O"

- name: 3-hydroxypipecolic acid
  smiles: "C1CC(C(NC1)C(=O)O)O"

- name: 3-hydroxyproline
  smiles: "OC1C(NCC1)C(=O)O"

- name: 3-methylproline
  smiles: "CC1C(NCC1)C(=O)O"

- name: 4,5-dehydroarginine
  smiles: "O=C(O)C(N)CC=CNC(N)=N"

- name: 4,5-dihydroxyornithine
  smiles: "C(C(C(=O)O)N)C(C(N)O)O"

- name: 4-acetamidopyrrole-2-carboxylic acid
  smiles: "CC(=O)NC1=CNC(=C1)C(=O)O"

- name: 4-amino-2-hydroxy-3-isopropoxybenzoic acid
  smiles: "CC(C)OC1=C(C=CC(=C1O)C(=O)O)N"

- name: 4-aminobutyric acid
  smiles: "NCCCC(=O)O"

- name: 4-aminophenylalanine
  smiles: "C1=CC(=CC=C1CC(C(=O)O)N)N"

- name: 4-chlorobenzoic acid
  smiles: "C1=CC(=CC=C1C(=O)O)Cl"

- name: 4-hydroxy-3-nitrobenzoic acid
  smiles: "C1=CC(=C(C=C1C(=O)O)[N+](=O)[O-])O"

- name: 4-hydroxy-D-kynurenine
  smiles: "C1=C(O)C=C(C(=C1)C(=O)CC(C(=O)O)N)N"

- name: 4-hydroxybenzoic acid
  smiles: "C1=CC(=CC=C1C(=O)O)O"

- name: 4-hydroxyglutamine
  smiles: "C(C(O)C(=O)N)C(C(=O)O)N"

- name: 4-hydroxyindole-3-carboxylic acid
  smiles: "c1cc2c(c(c1)O)c(c[nH]2)C(=O)O"

- name: 4-hydroxyphenylpyruvic acid
  smiles: "C1=CC(=CC=C1CC(=O)C(=O)O)O"

- name: 4-hydroxythreonine
  smiles: "C(C(C(C(=O)O)N)O)O"

- name: 4-hydroxyvaline
  smiles: "CC(CO)C(C(=O)O)N"

- name: 4-methoxytryptophan
  smiles: "C1=CC=C2C(=C1OC)C(=CN2)CC(C(=O)O)N"

- name: 4-nitrotryptophan
  smiles: "C1=CC=C2C(=C1[N+](=O)[O-])C(=CN2)CC(C(=O)O)N"

- name: 4-oxoproline
  smiles: "C1C(NCC1=O)C(=O)O"

- name: 4-butenyl-4-methylthreonine
  smiles: "CC=CCC(C)C(C(C(=O)O)N)O"

- name: 4-hydroxyproline
  smiles: "C1C(NCC1O)C(=O)O"

- name: 4-propylproline
  smiles: "CCCC1CC(NC1)C(=O)O"

- name: 4,5-dihydroxy-2-aminopentanoic acid
  smiles: "OC(CC(C(=O)O)N)CO"

- name: 4-acetyl-5-methylproline
  smiles: "CC(=O)OC1CC(NC(C)1)C(=O)O"

- name: 4-hydroxylysine
  smiles: "NCCC(O)CC(C(=O)O)N"

- name: 4-methylazetidine-2-carboxylic acid
  smiles: "CC1CC(N1)C(=O)O"

- name: 4-methylproline
  smiles: "CC1CC(NC1)C(=O)O"

- name: 4-propenylproline
  smiles: "CC=CC1CC(NC1)C(=O)O"

- name: 5-hydroxylysine
  smiles: "NCC(CCC(C(O)=O)N)O"

- name: 5,5-dimethylpipecolic acid
  smiles: "C1C(C)(C)CNC(C1)C(=O)O"

- name: 5-aminolevulinic acid
  smiles: "C(CC(=O)O)C(=O)CN"

- name: 5-chloroanthranilic acid
  smiles: "C1=CC(=C(C=C1Cl)C(=O)O)N"

- name: 5-chlorotryptophan
  smiles: "C1=CC2=C(C=C1Cl)C(=CN2)CC(C(=O)O)N"

- name: 5-methoxytyrosine
  smiles: "C1=C(OC)C(=CC=C1CC(C(=O)O)N)O"

- name: 5-methylorsellinic acid
  smiles: "C=1(C=C(C(=C(C1C)C)C(=O)O)O)O"

- name: 5-methylproline
  smiles: "C1CC(NC(C)1)C(=O)O"

- name: 6,7-dichlorotryptophan
  smiles: "C1=C(Cl)C(Cl)=C2C(=C1)C(=CN2)CC(C(=O)O)N"

- name: 6-chloro-4-hydroxy-1-methyl-indole-3-carboxylic acid
  smiles: "C(O)1=C(Cl)C=C2C(=C1)C(=CN(C)2)C(=O)O"

- name: 6-chloro-4-hydroxyindole-3-carboxylic acid
  smiles: "c(Cl)1cc2c(c(c1)O)c(c[nH]2)C(=O)O"

- name: 6-chlorotryptophan
  smiles: "C1=C(Cl)C=C2C(=C1)C(=CN2)CC(C(=O)O)N"

- name: 6-hydroxy-tetrahydro-isoquinoline-3-carboxylic acid
  smiles: "C1C(NCC2=C1C=C(C=C2)O)C(=O)O"

- name: 6-methyl-pipecolic acid
  smiles: "C1CC(C)NC(C1)C(=O)O"

- name: An acid hydrazine polyene (intermediate 14)
  smiles: "OC(=O)CCC(=O)NNCC(=O)O"

- name: Compound 4 (formed by the decarboxylative condensation of L-Phe and succinyl-CoA)
  smiles: "C1=CC=C(C=C1)CC(C(=O)CCC(=O)O)N"

- name: isovaline
  smiles: "CCC(C)(C(=O)O)N"

- name: lysergic acid
  smiles: "CN1CC(C=C2C1CC3=CNC4=CC=CC2=C34)C(=O)O"

- name: N-(1-methyl)-tryptophan
  smiles: "C1=CC=C2C(=C1)C(=CN(C)2)CC(C(=O)O)N"

- name: N-(1-propargyl)-tryptophan
  smiles: "C1=CC=C2C(=C1)C(=CN(CC#C)2)CC(C(=O)O)N"

- name: N-formylglycine
  smiles: "C(C(=O)O)NC=O"

- name: N-hydroxyvaline
  smiles: "CC(C)C(C(=O)O)NO"

- name: N-methylphenylalanine
  smiles: "CNC(CC1=CC=CC=C1)C(=O)O"

- name: N-methyltyrosine
  smiles: "C1=CC(=CC=C1CC(C(=O)O)NC)O"

- name: N1-methoxytryptophan
  smiles: "C1=CC=C2C(=C1)C(=CN(OC)2)CC(C(=O)O)N"

- name: N5-acetyl-hydroxyornithine
  smiles: "CC(=O)N(CCCC(C(=O)O)N)O"

- name: N5-nitroso-N5-hydroxyornithine
  smiles: "O=NN(CCCC(C(=O)O)N)O"

- name: N5-anhydromevalonyl-N5-hydroxyornithine
  smiles: "C(CC(C(=O)O)N)CN(O)C(=O)C=C(C)CCO"

- name: N6-hydroxylysine
  smiles: "C(CCNO)CC(C(=O)O)N"

- name: 3-hydroxy-3-methylproline
  smiles: "OC(C)1C(NCC1)C(=O)O"

- name: aza-beta-tyrosine
  smiles: "C1=CC(=NC=C1O)C(CC(=O)O)N"

- name: beta-methyltryptophan
  smiles: "CC(C1=CNC2=CC=CC=C21)C(C(=O)O)N"

- name: beta-phenylalanine
  smiles: "C1=CC=C(C=C1)C(CC(=O)O)N"

- name: adenosylmethionine
  smiles: "C[S+](CCC(C(=O)[O-])N)CC1C(C(C(O1)N2C=NC3=C(N=CN=C32)N)O)O"

- name: beta-hydroxycyclohex-2-enylalanine
  smiles: "C1C=CC(CC1)C(O)C(C(=O)O)N"

- name: beta-hydroxyenduracididine
  smiles: "C1C(NC(=N1)N)C(O)C(C(=O)O)N"

- name: beta-methylphenylalanine
  smiles: "CC(C1=CC=CC=C1)C(C(=O)O)N"

- name: beta-tyrosine
  smiles: "C1=CC(=CC=C1C(CC(=O)O)N)O"

- name: alaninol
  smiles: "CC(CO)N"

- name: argininol
  smiles: "NC(CO)CCCN=C(N)N"

- name: azetidine-2-carboxylic acid
  smiles: "O=C(O)C1NCC1"

- name: benzoxazolinate
  smiles: "c1ccc2c(c1)nc(o2)C(=O)O"

- name: beta-hydroxy-3-hydroxy-O-methyl-5-methyltyrosine
  smiles: "C1=C(C)C(=C(O)C=C1C(O)C(C(=O)O)N)OC"

- name: beta-hydroxy-gamma-methyl-hexadecanoic acid
  smiles: "CCCCCCCCCCCCC(C)C(O)CC(=O)O"

- name: beta-hydroxyarginine
  smiles: "C(C(O)C(C(=O)O)N)CN=C(N)N"

- name: beta-hydroxyphenylalanine
  smiles: "OC(C1=CC=CC=C1)C(C(=O)O)N"

- name: beta-lysine
  smiles: "C(CC(CC(=O)O)N)CN"

- name: betaine
  smiles: "C[N+](C)(C)CC(=O)O"

- name: caffeic acid
  smiles: "OC(=O)C=Cc1ccc(O)c(O)c1"

- name: capreomycidine
  smiles: "C1CN=C(NC1C(C(=O)O)N)N"

- name: cinnamic acid
  smiles: "C1=CC=C(C=C1)C=CC(=O)O"

- name: citrulline
  smiles: "C(CC(C(=O)O)N)CNC(=O)N"

- name: colletorin D acid
  smiles: "CC1=CC(=C(C(=C1C(=O)O)O)CC=C(C)C)O"

- name: coumaric acid
  smiles: "C1=CC(=CC=C1C=CC(=O)O)O"

- name: cysteic acid
  smiles: "C(C(C(=O)O)N)S(=O)(=O)O"

- name: dehydroarginine
  smiles: "C(CN=C(N)N)C=C(C(=O)O)N"

- name: dehydrophenylalanine
  smiles: "NC(=CC1=CC=CC=C1)C(=O)O"

- name: dehydrotryptophan
  smiles: "C1=CC=C2C(=C1)C(=CN2)C=C(C(=O)O)N"

- name: dehydrovaline
  smiles: "CC(=C(C(=O)O)N)C"

- name: dehydroalanine
  smiles: "C=C(N)C(=O)O"

- name: dihydrolysergic acid
  smiles: "CN1CC(CC2C1CC3=CNC4=CC=CC2=C34)C(=O)O"

- name: dimethylsulfoniopropionic acid
  smiles: "C[S+](C)CCC(=O)O"

- name: enduracididine
  smiles: "C1C(NC(=N1)N)CC(C(=O)O)N"

- name: fumaric acid
  smiles: "C(=CC(=O)O)C(=O)O"

- name: glycolic acid
  smiles: "C(C(=O)O)O"

- name: grifolic acid
  smiles: "CC(C)=CCCC(C)=CCCC(C)=CCC1=C(O)C=C(C)C(C(=O)O)=C(O)1"

- name: homophenylalanine
  smiles: "C1=CC=C(C=C1)CCC(C(=O)O)N"

- name: homoserine
  smiles: "C(CO)C(C(=O)O)N"

- name: homotyrosine
  smiles: "C1=CC(=CC=C1CCC(C(=O)O)N)O"

- name: homocysteine
  smiles: "NC(CCS)C(=O)O"

- name: indole pyruvic acid
  smiles: "C1=CC=C2C(=C1)C(=CN2)CC(=O)C(=O)O"

- name: leucinol
  smiles: "CC(C)CC(CO)N"

- name: linoleic acid
  smiles: "CCCCCC=CCC=CCCCCCCCC(=O)O"

- name: malonic acid
  smiles: "O=C(O)CC(=O)O"

- name: malic acid
  smiles: "C(C(C(=O)O)O)C(=O)O"

- name: malonamate
  smiles: "NC(=O)CC(=O)O"

- name: meta-tyrosine
  smiles: "C1=CC(=CC(=C1)O)CC(C(=O)O)N"

- name: methylglutaconyl hydroxyornithine
  smiles: "CC(=CC(=O)N(CCCC(C(=O)O)N)O)CC(=O)O"

- name: nicotinic acid
  smiles: "C1=CC(=CN=C1)C(=O)O"

- name: norcoronamic acid
  smiles: "CC1CC1(C(=O)O)N"

- name: ochratoxin beta
  smiles: "CC1CC2=C(C(=C(C=C2)C(=O)O)O)C(=O)O1"

- name: p-hydroxybenzoylformic acid
  smiles: "C1=CC(=CC=C1C(=O)C(=O)O)O"

- name: p-hydroxymandelate
  smiles: "C1=CC(=CC=C1C(C(=O)O)O)O"

- name: para-aminobenzoic acid
  smiles: "O=C(O)c1ccc(N)cc1"

- name: phenazine-1,6-dicarboxylic acid
  smiles: "C1=CC(=C2C(=C1)N=C3C(=N2)C=CC=C3C(=O)O)C(=O)O"

- name: phenylalaninol
  smiles: "C1=CC=C(C=C1)CC(CO)N"

- name: phenylglycine
  smiles: "C1=CC=C(C=C1)C(C(=O)O)N"

- name: phenyllactic acid
  smiles: "C1=CC=C(C=C1)CC(C(=O)O)O"

- name: phenylpyruvic acid
  smiles: "C1=CC=C(C=C1)CC(=O)C(=O)O"

- name: piperazic acid
  smiles: "C1CC(NNC1)C(=O)O"

- name: piperonylic acid
  smiles: "OC(=O)c1ccc2OCOc2c1"

- name: pyrrole-2-carboxylic acid
  smiles: "C1=CNC(=C1)C(=O)O"

- name: pyruvic acid
  smiles: "CC(=O)C(=O)O"

- name: quinoxaline-2-carboxylic acid
  smiles: "C1=CC=C2C(=C1)N=CC(=N2)C(=O)O"

- name: succinic semialdehyde
  smiles: "C(CC(=O)O)C=O"

- name: succinyl-hydrazinoacetic acid
  smiles: "NN=CC=CC=CC=CC=CC=CC(=O)O"

- name: tricarballylic acid
  smiles: "C(C(CC(=O)O)C(=O)O)C(=O)O"

- name: ustethylinic acid
  smiles: "c1(C)c(O)c(C(=O)O)c(CC)cc(O)1"

- name: valine isocyanide
  smiles: "CC(C)C([N+]#[C-])C(O)=O"

- name: valinol
  smiles: "CC(C)C(CO)N"

- name: parahydroxy phenylacetic acid
  smiles: "O=C(O)Cc1ccc(O)cc1"

- name: tryptophan alkaloid
  smiles: "O=C(O)C(CC12O)N[C@@H]2Nc3c1cccc3"

- name: guanidine-containing fatty acid
  smiles: "N=C(NCCCCCC(O)=O)N"

- name: guanidine-containing fatty acid
  smiles: "N=C(NCCCC=CC(O)=O)N"

- name: guanidine-containing fatty acid
  smiles: "N=C(N)NCCCC(=O)O"

- name: guanidine-containing fatty acid
  smiles: "N=C(N)NCC(=O)O"

- name: isoleucinol
  smiles: "CCC(C(N)CO)C"

- name: putrescin
  smiles: "NCCCCN"

- name: cadaverine
  smiles: "NCCCCCN"

- name: N-(5-aminopentyl)hydroxylamine
  smiles: "NCCCCCNO"
  pseudonyms: ["siderophore"]

- name: butanedioic acid
  smiles: "O=C(O)CCC(O)=O"
  pseudonyms: ["siderophore"]

- name: 3-formamido-2-hydroxybenzoic acid
  smiles: "O=CNc1cccc(C(=O)O)c1O"

- name: 2-phenylethanamine
  smiles: "NCCc1ccccc1"

- name: homocysteamine
  smiles: "NCCCS"

- name: valienol
  smiles: "OC1C=C(C(C(C1O)O)O)CO"

- name: valienone
  smiles: "OC1C=C(C(C(C1O)O)=O)CO"

- name: streptamine
  smiles: "NC1CC(C(C(C1O)O)O)N"
```
