## Supplementary material for "RetroMol: Parsing a shared encoding from natural products and their biosynthetic gene clusters": RetroMol Supplemental Information: Supplemental_Information.pdf

### S1: Full polyketide encoding scheme

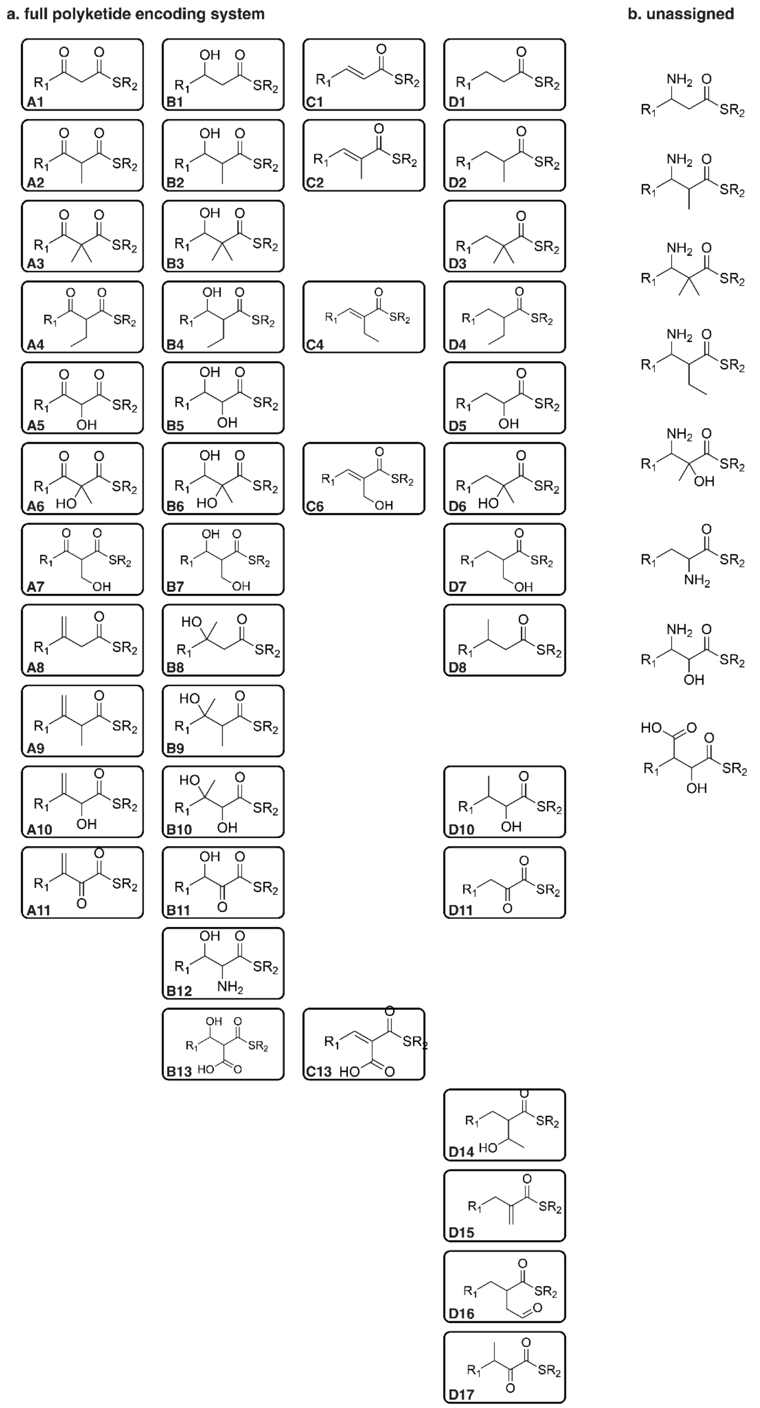

**S1**: Full polyketide encoding scheme. (a) Encoded polyketide building blocks found in retrosynthesized natural compounds from NPAtlas. Non-polyketide building blocks are encoded by their given name in the matching rule set, and abbreviated using the first three unique set of alphanumeric characters from their name. (b) Found but unencoded polyketide-like building blocks found in retrosynthesized natural compounds from NPAtlas.

### S2: Metabolomics analysis for nocardichelin B deorphanization

| *Nocardia* sp. NBC | 00511 | 01009 | 01329 | 01730 |
|---|---|---|---|---|
| 716.459 *m/z* | ⊠ | ⊠ | ⊠ | ⊠ |
| Candidate BGC | ⊠ | ⊠ | ⊠ | ⊠ |

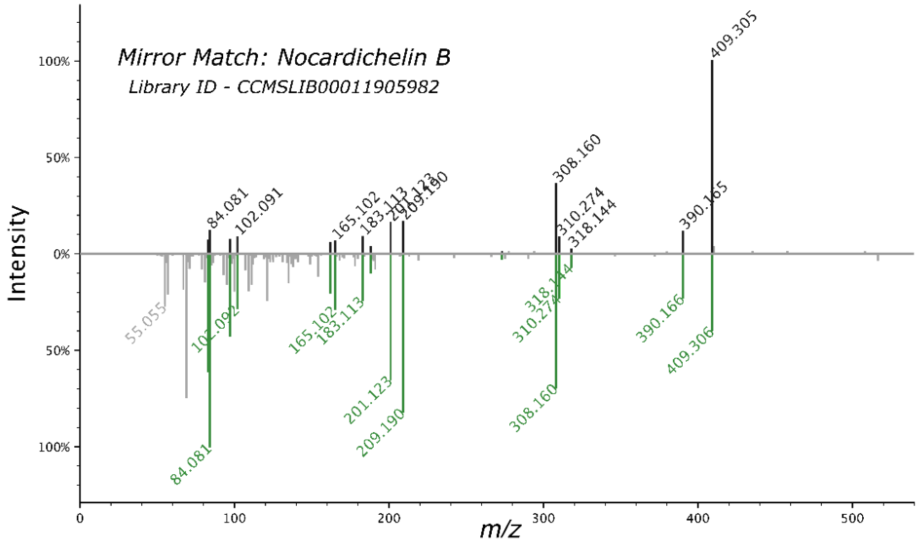

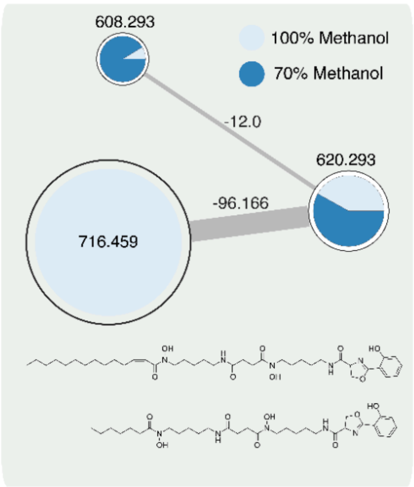

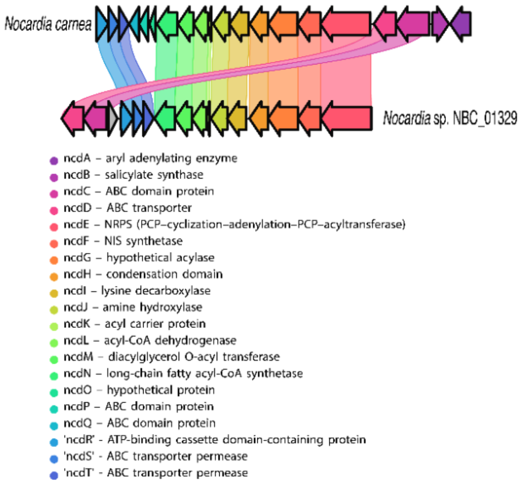

**S2**: Metabolomics analysis for the deorphanization of nocardichelin B.

### S3: Example reaction and assembly graphs for various input structures

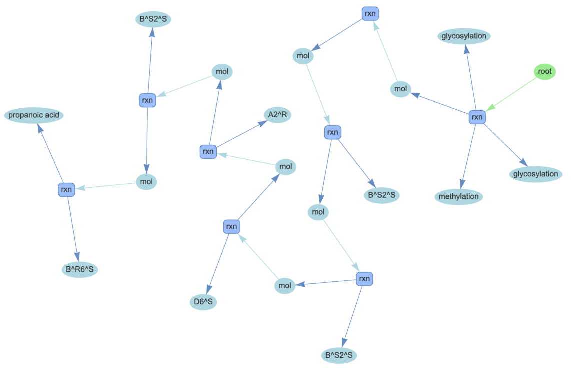

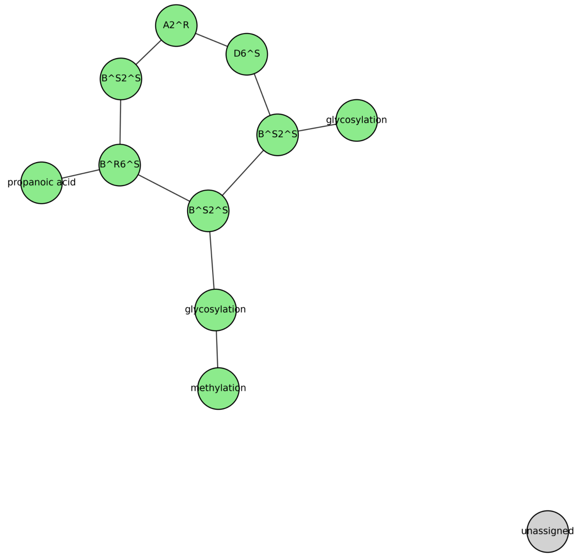

**S3a**: reaction graph for erythromycin C.

**S3b**: assembly graph for erythromycin C.

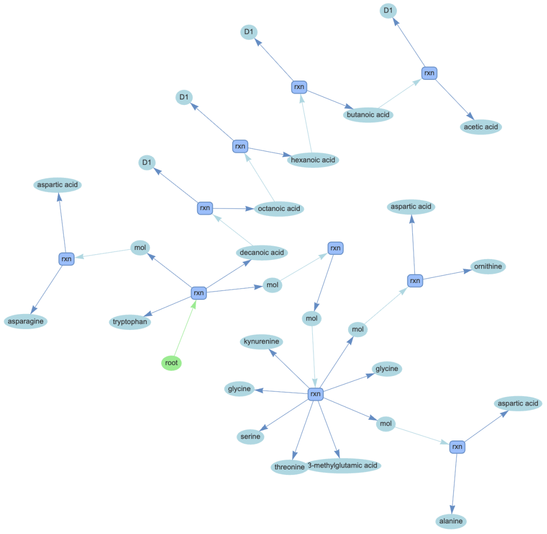

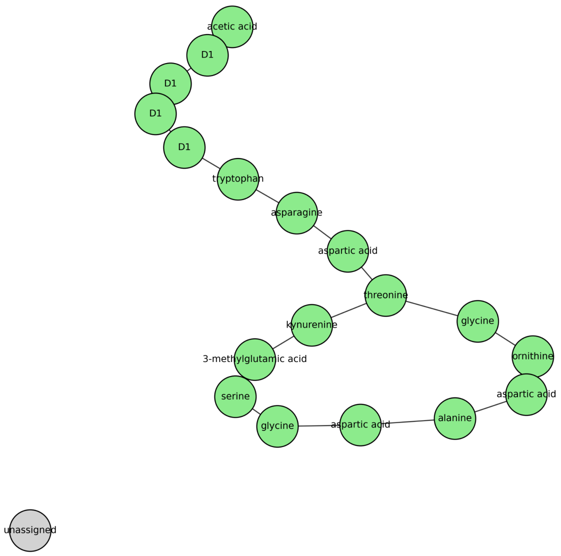

**S3c**: reaction graph for daptomycin.

**S3d**: assembly graph for daptomycin.

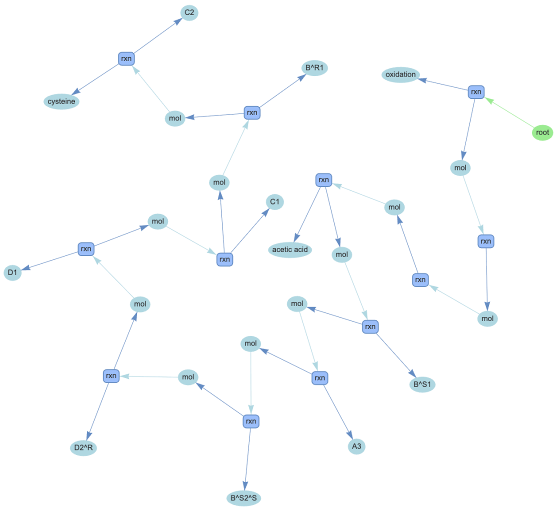

**S3e**: reaction graph for epothilone.

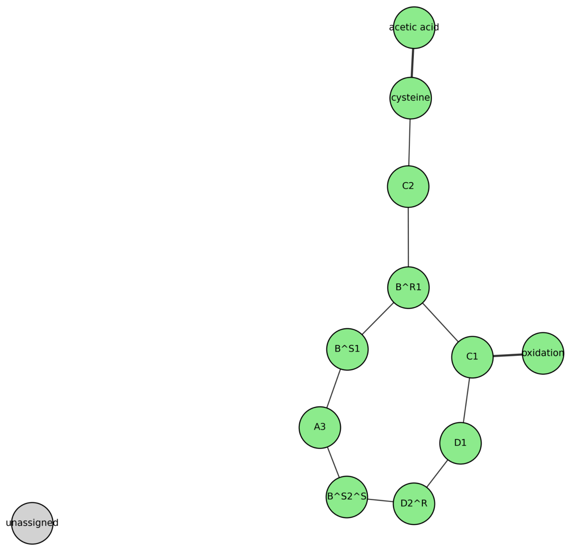

**S3f**: assembly graph for epothilone.
