## Supplementary figures and images for "RetroMol: Parsing a shared encoding from natural products and their biosynthetic gene clusters"

### assembly_graph.png

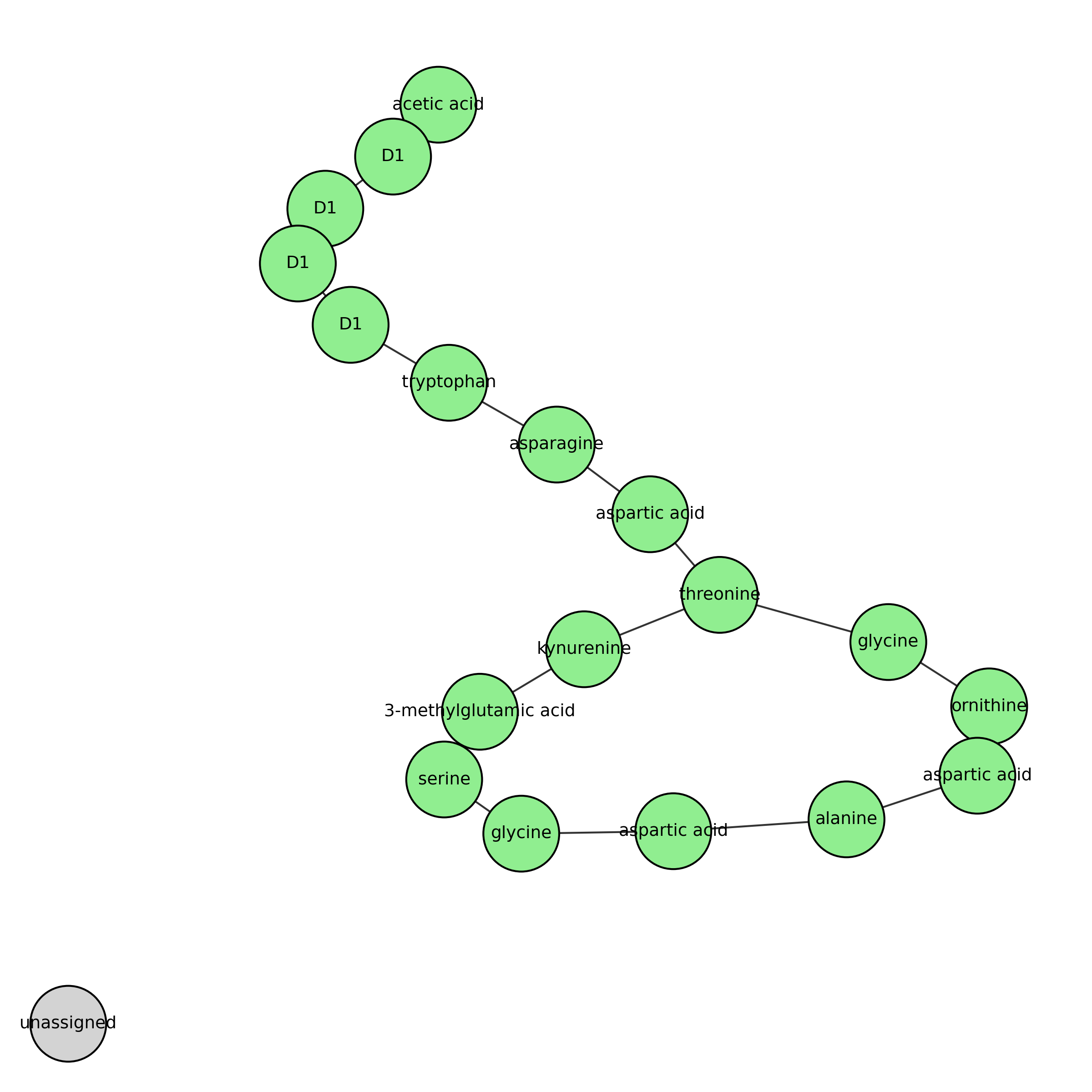

### assembly_graph.png

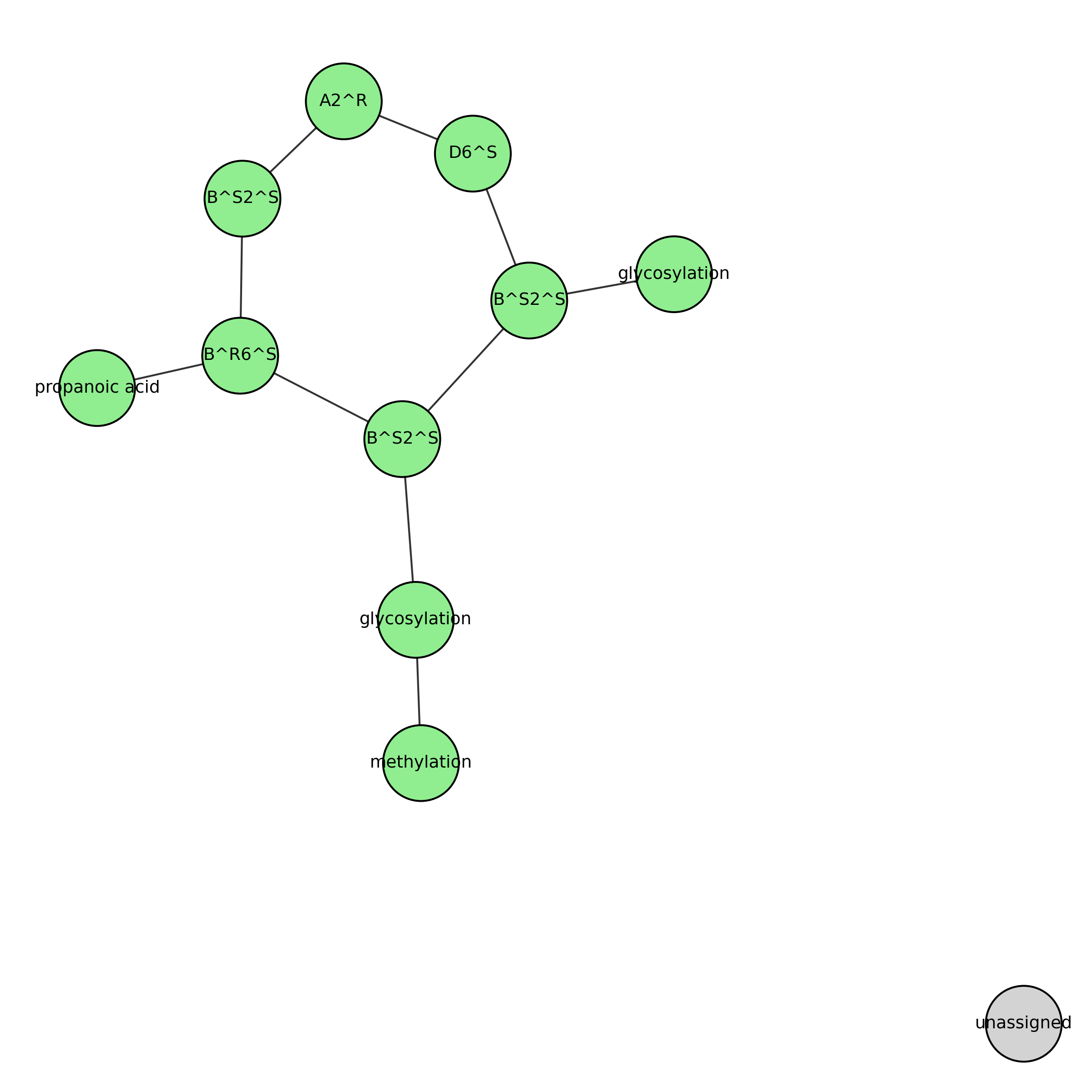

### assembly_graph.png

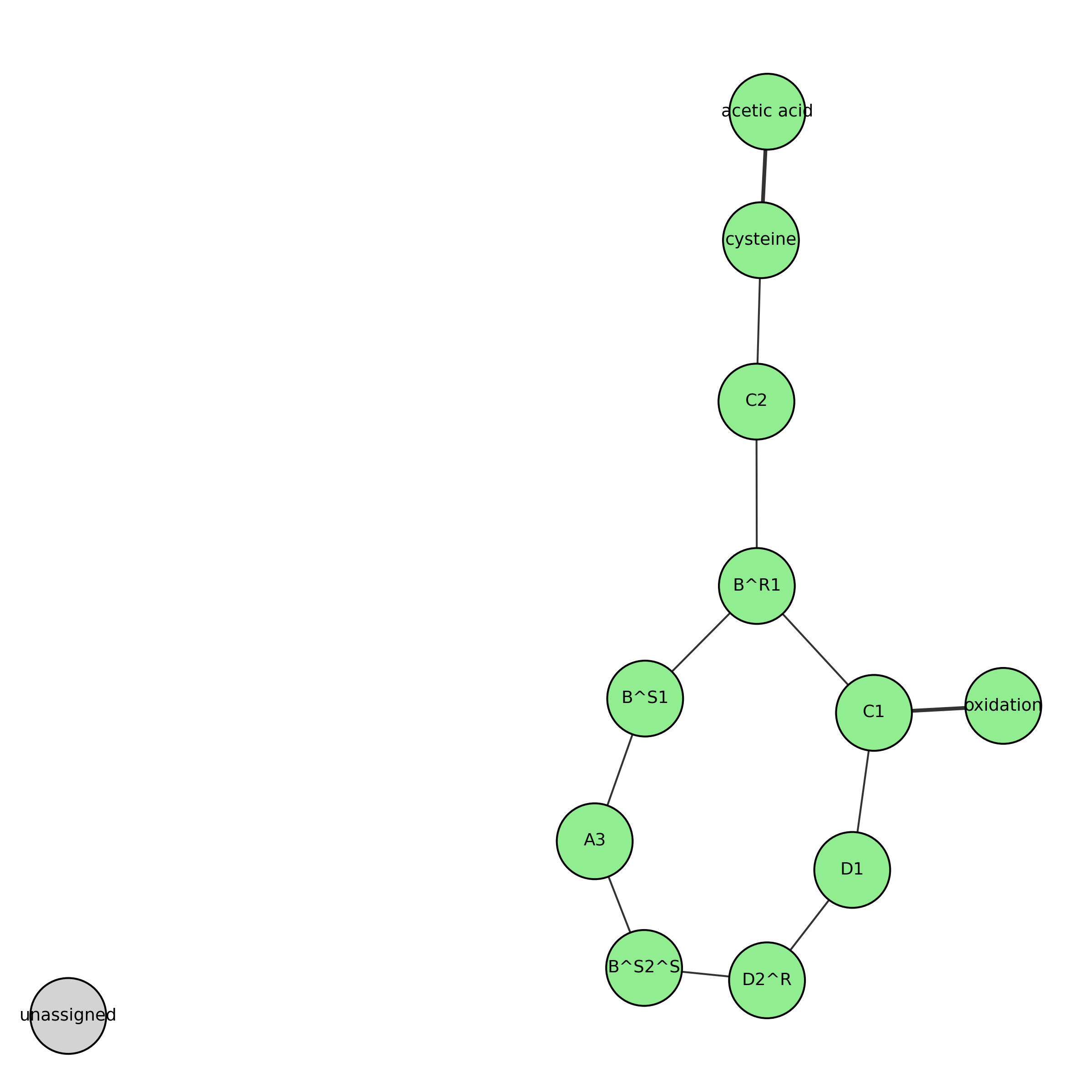
